# Biology of Coital Behavior: Looking Through the Lens of Mathematical Genomics

**DOI:** 10.1101/2023.04.12.536521

**Authors:** Moumita Sil, Debaleena Nawn, Sk. Sarif Hassan, Subhajit Chakraborty, Arunava Goswami, Pallab Basu, Lalith Roopesh, Emma Wu, Kenneth Lundstrom, Vladimir N. Uversky

**Affiliations:** Biological Science Division, Indian Statistical Institute, 203 B.T Road, Kolkata, 700108, West Bengal, India; Tata Medical Center, 14 MAR (E-W), New Town, Rajarhat, Kolkata, 700160, West Bengal, India; Department of Mathematics, Pingla Thana Mahavidyalaya, Maligram, Paschim Medinipur, 721140, West Bengal, India; School of Physics, University of the Witwatersrand, Johannesburg, Braamfontein 2000, South Africa; Department of Molecular Medicine, Morsani College of Medicine, University of South Florida, Tampa, FL 33612, USA; PanTherapeutics, Rte de Lavaux 49, CH1095 Lutry, Switzerland

**Author notes:** ^∗^*These authors contributed equally in the work (Moumita Sil, Debaleena Nawn, Sk. Sarif Hassan). Corresponding authors: Vladimir N. Uversky and Sk. Sarif Hassan Email addresses (Moumita Sil), (Debaleena Nawn), (Sk. Sarif Hassan), (Subhajit Chakraborty), (Arunava Goswami), (Pallab Basu), (Lalith Roopesh), (Emma Wu), (Kenneth Lundstrom), (Vladimir N. Uversky).

**Keywords:** Vasopressin, Dopamine, Oxytocin, Monogamy-Polygamy, Coital behavior, Proximal relationship.

## Abstract

Research has shown that genetics and epigenetics regulate mating behavior across multiple species. Previous studies have generally focused on the signaling pathways involved and spatial distribution of the associated receptors. However a thorough quantitative characterization of the receptors involved may offer deeper insight into mating behavioral patterns. Here oxytocin, arginine-vasopressin 1a, dopamine 1, and dopamine 2 receptors were investigated across 76 vertebrate species. The receptor sequences were characterized by polarity-based randomness, amino acid frequency-based Shannon entropy and Shannon sequence variability, intrinsic protein disorder, binding affinity, stability and pathogenicity of homology-based SNPs, structural and physicochemical features. Hierarchical clustering of species was derived based on structural and physicochemical features of the four receptor sequences separately, which eventually led to proximal relationships among 29 species. Humans were found to be significantly distant phylogenetically from the prairie voles, a representative of monogamous species based on coital behavior. Furthermore, the mouse (polygamous), the prairie deer mouse (polygamous), and the prairie vole (monogamous) although being proximally related (based on quantitative genomics of receptors), differed in their coital behavioral pattern, mostly, due to behavioral epigenetic regulations. This study adds a perspective that receptor genomics does not directly translate to behavioral patterns.

## 1. Introduction

Genes play an important role in controlling the underlying mechanisms and neurochemistry which are responsible for the social behavior of different animals, including humans. In terms of mate selection and bond-formation with mates, the animal kingdom in general exhibits two kinds of behavior: monogamous and polygamous [1]. In a monogamous mating system, each sex has a single sexual partner for a particular period of time [2]. Monogamy generally appears to be low in the animal kingdom, but is mostly prevalent in birds, while very few mammals and even lesser number of invertebrates exhibit monogamous mating systems [3]. However, the so called “monogamous” species are quite variable in terms of their social and reproductive behaviours [4]. This has given rise to separate terms such as “social monogamy” and “reproductive” or “genetic monogamy”. In social monogamy, the male and female show behaviours such as shared use of territory, bi-parental care and other behavior indications of being a social couple, but do not necessarily copulate and reproduce with only one partner [3]. Social monogamy is widely observed in birds where more than 90% of the species are socially monogamous [5]. In mammals, only about 9% of all species have been identified as socially monogamous [4]. However, as mentioned before, social monogamy does not necessarily imply an exclusive sexual relationship between males and females. For example, in the prairie voles (*Microtus ochrogaster*), which has been presented as a model organism for social monogamy, can also have additional partners in the wild [6]. Only a few sexually reproducing organisms exhibit exclusive sexual lifetime monogamy (reproductive/genetic monogamy) [3]. Also, genetic monogamy may be observed between individual couples of socially monogamous species, where behaviours such as mate-guarding may prevent mating with other individuals [7, 8]. Polygamy on the other hand is the most prevalent mating behavior in the animal kingdom which involves mating with multiple individuals of the opposite sex. Polygamy exhibits several forms such as polyandry (an individual female mating with multiple males), polygyny (an individual male mating with multiple females) and polygyandry or promiscuity (mating between multiple members of both sexes) [9]. Such diversity of mating behavior among animals may be governed by the presence of hormones – oxytocin and arginine-vasopressin and the neurotransmitter dopamine. Both hormones and neurotransmitters have been shown to play a regulatory role in pair-bond formation among the socially monogamous prairie vole (*Microtus ochrogaster*) [10, 11, 12]. In prairie voles, the action of arginine-vasopressin neuropeptide on the vasopressin-1a receptor (V1aR) has been shown to regulate different types of social behavior, including social bonding between males and females [13, 14]. Several studies have also shown that the concentration and spatial distribution of V1aR may correlate with inter-specific and intra-specific differences in social attachment of male organisms for different species withe the Microtus genus [15]. In the socially monogamous prairie vole (*Microtus ochrogaster*), V1aR is expressed at greater concentration in the ventral forebrain as compared to the promiscuous meadow vole (*Microtus pennsylvanicus*) [16]. The gene encoding the V1a receptor, AVPR1a, regulates the distribution of V1aR in the forebrain and therefore plays an important role in the regulation of social attachment. Polymorphism in the regulatory 5’ region of the microsatellite DNA of the avpr1a gene is found in monogamous prairie voles, but not in promiscuous meadow voles [17]. By increasing the expression of AVPR1a gene through viral vector mediated gene transfer, promiscuous male meadow voles show selective pair-bonding with females in a manner similar to monogamous prairie voles [18].

Some studies have shown that central infusion of oxytocin increases social pair-bonding in prairie voles, whereas the administration of oxytocin antagonists inhibits such behavior [11, 19, 20]. The density of the oxytocin receptor OXTR has been found to be higher in the nucleus accumbens region of the brain in socially monogamous vole species as compared to non-monogamous voles [21]. Needless to say, the expression of the OXTR gene encoding the oxytocin receptor plays an important role in this regard. Over-expression of the OXTR gene through viral vector mediated gene transfer in pre-pubertal female prairie voles led to display of partner preference and alloparental care at the adult stage [22]. However, another study published in 2022 employed CRISPR to produce Oxtr-null mutants of prairie voles and these voles showed partner preference, aggression towards other potential mates and biparental care even in the absence of OXTR expression [23]. It has been hypothesized that oxytocin may also function through a different pathway to exhibit such behavior. Oxytocin has been known to signal via the AVPR1a gene as well, and therefore these two pathways may be redundant in regulating mate-selection and biparental care [24, 25]. The role of dopamine in regulation of social behavior has also been studied in socially monogamous prairie voles. The administration of dopamine antagonist haloperidol prevented partner preference in prairie voles, whereas the dopamine agonist apomorphine induced this behavior [26]. Among dopamine DRD1 and DRD2 receptors, DRD2 receptors have been shown to play an important role in social attachment in prairie voles [27, 28]. Studies have also shown that the expression of both the OXTR and DRD2 type receptors in the nucleus accumbens (NAcc) regulates the social behavior of prairie voles [29]. This socially monogamous rodent, Microtus ochrogaster has actually been used in numerous studies as an animal model for understanding the neurobiological basis of social attachment, monogamy and parental care and these studies transcend to human social behavior as well [30]. In the present study, we investigated the V1AR/AVPR1A, OXTR, and the DRD1 and DRD2 receptors from 76 vertebrate organisms and attempted to establish a proximal relationship among them based on different quantitative analyses of their genetic sequences. The relatedness and/or divergence between these sequences may help us to find a possible correlation with the variation of social behavior (monogamy or polygamy) among different species. In order to validate the proposed hypothesis, further research and hands-on experimentation may be required.

## 2. Data acquisition

In the present study, 117 AVPR1a, 83 DRD1, 174 DRD2, and 91 OXTR receptor sequences from 76 species with different frequencies were investigated. For all species, those with at least one sequence available from all four receptors were included (Table 1).

**Table 1:**
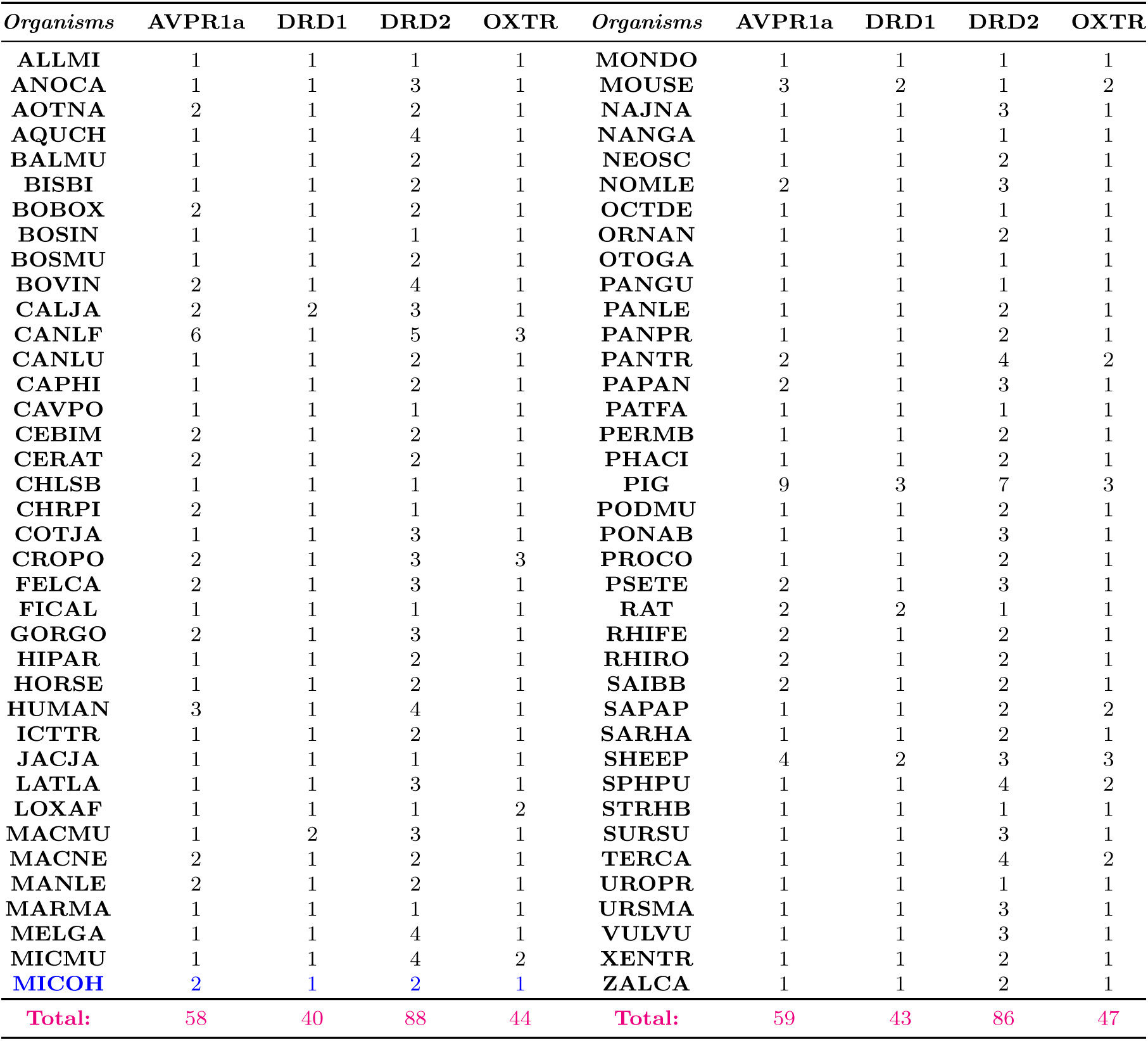
List of 76 species from which AVPR1a, DRD1, DRD2, and OXTR receptor sequences with respective frequencies were considered. In the list code names were used as per Unirpot code. The scientific names and their associated code names were listed in **Supplementary file-1**

List of all accession numbers of all receptor sequences was made available in the folder Sequence files. Furthermore, it was noted that the following pairs of sequences were 100%. identical **Supplementary file-2**.

The 76 species that were taken into account were all under the phylum Chordata. We had further grouped these organisms in terms of their class, order and coital behaviour (Figure 1). The mating behavioral pattern of these species were obtained from websites and articles Animal Diversity Web, Primate Info Net, [31, 32, 33, 34, 35, 36, 37, 38, 39, 40, 41, 42, 43, 44, 45, 46, 47, 48, 49, 50, 51, 52, 53, 54, 55, 56, 57, 58, 59, 60, 61, 62, 63, 64, 65, 66, 67, 68, 69, 64, 70, 71, 72, 73, 74, 75, 76, 77, 78, 79, 80, 81].

**Figure 1:**
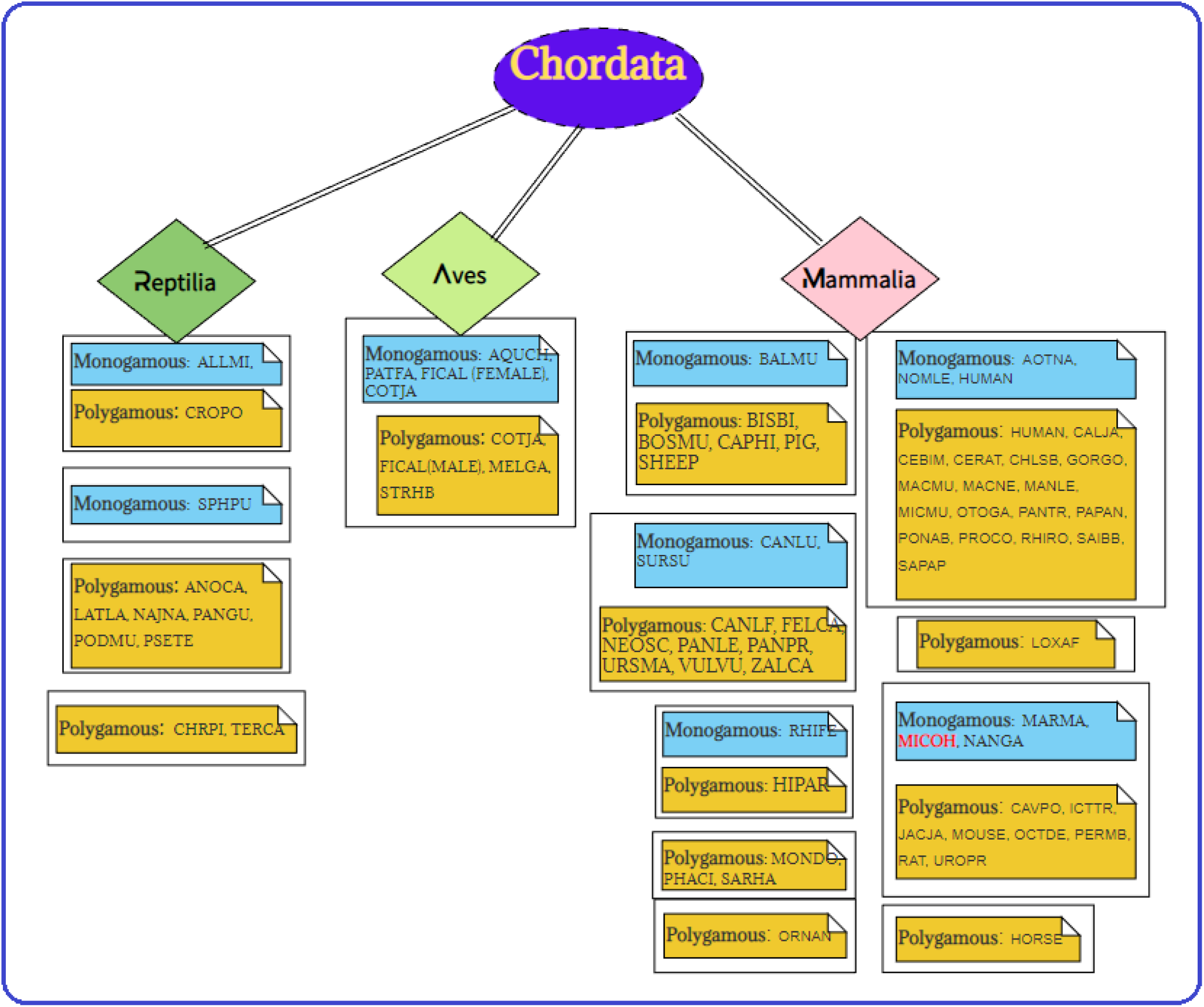
Monogamy-Polygamy behavior of the species as known from various literature as cited above (31-81)

## 3. Methods

### 3.1. Quantitative features extracted from AVPR1a, DRD1, DRD2, and OXTR receptors

#### 3.1.1. Deviation from randomness based on polarity

We studied the sequence of four neurotransmitter receptors AVPR1a, DRD1, DRD2, and OXTR. We wanted to investigate how amino acid sequences of the receptors deviate from randomness. A statistical deviation from randomness in the spatial arrangement of amino acids was studied by analyzing the clustering properties. First, every amino acid in a given receptor amino acid sequence was identified as polar(P) or non-polar(Q). Thus, every receptor amino acid sequence became a binary sequence with two symbols: P and Q. We tested the null hypothesis that such a string of P and Q is a random string with no-correlation between the placements of polar and non-polar amino acids. To test our null hypothesis we performed the following bootstrap analysis in three steps:

The first step of our analysis involved computing the average cluster size (m) of each binary sequence. Next, we randomly permuted each binary sequence 5000 times and recorded the mean (*µ*) and variance (*σ*^2^) of the resulting average cluster sizes. We defined *z* as being distributed according to 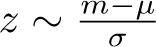, and we noted the value of *z*.

We performed the above two steps for all binary sequences. We checked our null hypothesis by performing a *Student-t-test* to check if the mean of *z* is significantly different from 0 [82, 83].

#### 3.1.2. Evaluating intrinsic protein disorder

Protein disorder refers to the absence of a well-defined three-dimensional structure of a protein or protein region under physiological conditions. Intrinsically disordered proteins (IDPs) are a subgroup of proteins that are particularly flexible and lack a stable, well-defined structure even under native conditions [84, 85]. Interestingly, they are still capable of carrying out varied biological functions.

To assess the per-residue disorder propensity within the amino acid sequences of AVPR1a, DRD1, DRD2, and OXTR receptors, we used PONDR® VSL2, which is a highly accurate standalone disorder predictor [86]. The per-residue disorder predisposition scores range from 0 to 1, where a score of 0 indicates fully ordered residues, and a score of 1 indicates fully disordered residues. Residues with scores above the threshold of 0.5 are considered *disordered residues*. Residues with disorder scores between 0.25 and 0.5 are categorized as *highly flexible*, while those with scores between 0.1 and 0.25 are classified as *moderately flexible* [86].

#### 3.1.3. Amino acid frequency based Shannon entropy of sequence

The Shannon entropy measures the information content of a system [87]. The formula for Shannon entropy of an amino acid sequence is:

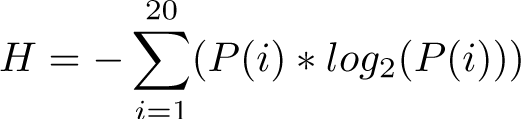

Where *H* is the Shannon entropy, *P* (*i*) is the frequency probability of occurrence of amino acid *i*, and the summation is over all possible amino acids [83]. The resulting value of *H* reflects the degree of diversity of frequency distribution of amino acids. A higher value of *H* indicates greater diversity, while a lower value indicates less diversity.

#### 3.1.4. Amino acid frequency based Shannon variability of position

Shannon entropy is deployed to estimate variability of amino acid residues at each residue position across all aligned sequences. The Shannon variability (*H_v_*) for every position is defined as follow:

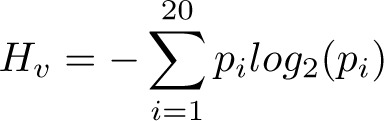

Here *p_i_* represents the fraction of residues of amino acid type *i* at a particular position. *H_v_* ranges from 0 (only one residue present at that position) to 4.322 (all 20 residues are equally represented in that position) [88]. Typically, positions with *H_v_ >* 2.0 are considered *variable*, whereas those with *H_v_ <* 2 are considered *conserved*. *Highly conserved* positions are those with *H_v_ <* 1.0 [88].

By analyzing per-residue variability using Shannon entropy, one can identify functionally important residues within the receptor protein family. For example, highly variable residues may be involved in ligand binding, while highly conserved residues may be critical for maintaining the overall structure and stability of the receptor. In addition, per-residue variability analysis using Shannon entropy can help identify potential drug targets within the receptor protein family. Residues that are highly variable among receptor sequences may be more amenable to small molecule binding, as they may have more flexibility and plasticity in their binding pockets [89].

#### 3.1.5. Binding affinity of receptors

Binding affinity alludes to the relative strength of the receptor-ligand binding by computationally calculating the conformational energy of the receptor and ligand in their strongest found orientation. Homology models of each receptor were created based on Uniprot sequences and the closest crystallographic structure was found using SWISS-MODEL. Global Model Quality Estimate (GMQE) and template used were recorded for each receptor. GMQE measures how well (1) the template matches the homology model and (2) the homology model matches the predicted tertiary structure based on the primary sequence [90, 91].

Models were prepared and docked using Autodock Tools and Autodock Vina in a SLURM system, and a protocol was created to ensure consistency between all runs. The strongest affinity (lowest energy) configuration was recorded as a basis of analysis. Extremely far outliers and errors in docking (resulting in null or unreasonable data) were found and redocked.

#### 3.1.6. Determining homology based SNPs, pathogenicity, and structural stability

To assess the single nucleotide polymorphism (SNP) across the remaining sequences for AVPR1a, DRD1, DRD2, and OXTR receptors, we used sp—Q9WTV9—V1AR MICOH, tr—E0V8E1—E0V8E1 MICOH D(1A), tr—E0V889—E0V889 MICOH D(2), and tr—E0V872—E0V872 MICOH as reference sequences, respectively. We used the Clustal Omega web-server to align the remaining sequences to the reference sequences and to identify similar non-synonymous changes in the amino acid residues, which we referred to as ”homology based SNPs” (in short simply ”SNPs”) [83].

Computational tools that predict the effects of mutations on protein function are crucial for analyzing single nucleotide variants and prioritizing them for experimental characterization. We used the PREDICTSNP web-server, which employs six reliable tools (MAPP, nsSNPAnalyzer, PANTHER, PhD-SNP, PolyPhen-1, PolyPhen-2, SIFT, and SNAP) to predict the pathogenicity of a given mutation [92]. The consensus prediction for mutations was robust and accurate, and we used this web-server to predict the pathogenicity (neutral or disease) of the SNPs detected for all receptor sequences.

A single mutation on an amino acid residue can cause a severe change in protein structure, disrupting its function. Predicting protein stability changes can help identify possible candidates for novel protein design. We used the iStable web-server, which uses either the protein structure or sequence as input [93]. In our study, we used the receptor protein sequences as input to evaluate the stability of each homology based SNP.

#### 3.1.7. Structural and physicochemical features (I-features and P-features)

Structural and physicochemical descriptors extracted from sequence data have been widely used to characterize sequences and predict structural, functional, expression and interaction profiles of proteins. iFeature (I-features), a versatile Python-based toolkit for generating various numerical feature representation schemes for protein sequences was deployed to extract structural and physicochemical properties of the four receptors [94].

A list of 1182 features which were deployed in the present study was summarized in Table 2.

**Table 2:** List of features of dimension 1182 calculated by I-feature for all receptor sequences

| List of various features (1182 dimensional) calculated by iFeature |  |  |
| --- | --- | --- |
| Descriptor groups | Descriptor | Dimension |
| Amino acid composition | Amino acid composition (AAC) | 20 |
|  | Dipeptide composition (DPC) | 400 |
| Grouped amino acid composition | Grouped Amino Acid Composition (GAAC) | 5 |
|  | Grouped Dipeptide Composition (GDC) | 25 |
| Autocorrelation | Moran Autocorrelation | 240 |
| C/T/D | C/T/D Composition (CTDC) | 39 |
| Conjoint Triad | Conjoint Triad | 343 |
| Quasi-sequence-order | Sequence-Order-Coupling Number (SoC number) | 60 |
| Pseudo-amino acid composition | Pseudo-Amino Acid Composition (PAAC) | 50 |

Furthermore, P-features is a web-server that provides a wide range of tools for computing various protein features. With its composition-based module, one can compute features such as: (i) tripeptide amino acid compositions; (ii) Repeats and distribution of amino acids; and (iii) Miscellaneous compositions, such as pseudo amino acid, autocorrelation, conjoint triad, and quasi-sequence order [95].

#### 3.1.8. Hierarchical clustering of sequences based on I-feature and P-feature

UPGMA (unweighed pair group method with arithmetic mean), an agglomerative hierarchical clustering method was deployed to cluster receptor sequences based on distance matrix (Euclidean metric) formed by I-features and P-features, separately for each of the four receptors [96].

## 4. Results and analyses

### 4.1. Deviation from randomness

We saw a huge disparity of randomness among AVPR1a, DRD1, DRD2, and OXTR (Figure 2). The *p−*value obtained for the receptors follows ascending order with respect to Mean(*z*),

- For DRD1 receptors, *z* = 0.50, Ttest 1sampResult(statistic=10.53, p-value= 6.82 *×* 10*^−^*^17^).
- For OXTR receptors, *z* = 0.52, Ttest 1sampResult(statistic=8.63, p-value= 1.99 *×* 10*^−^*^13^)
- For AVPR1a receptors, *z* = 1.26, Ttest 1sampResult(statistic=25.97, p-value= 3.62 *×* 10*^−^*^50^)
- For DRD2 receptors, *z* = 2.12, Ttest 1sampResult(statistic=42.96, p-value= 3.22 *×* 10*^−^*^94^)

**Figure 2:**
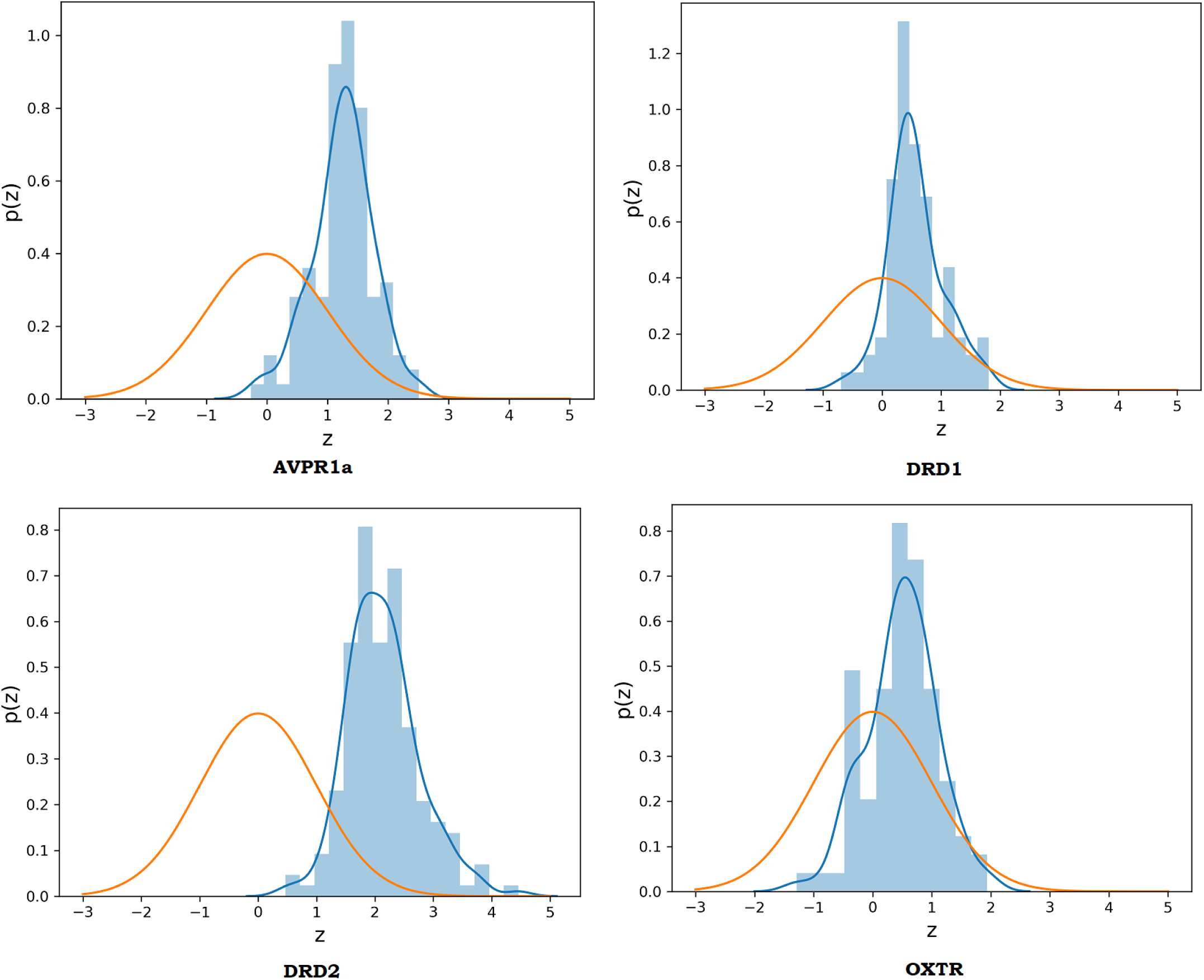
Difference between the standard normal and the distribution of average cluster sizes of polar-nonpolar sequences of AVPR1a (Top-left) and DRD1 (Top-right), DRD2 (Bottom-left), and OXTR (Bottom-right) receptors. Orange: standard normal, Blue: the histogram and kernel density of mean cluster sizes.

Hence, in all cases the null hypothesis was rejected. We noticed the highest deviation in the case of DRD2 and the smallest in DRD1. Furthermore, species level deviation in randomness by limiting our analysis to sequences belonging to a particular organism was carried out. For the DRD2 humans have a Mean(*z*) of 1.22 (PHACI shows the most deviation from randomness), which is lowest among all the species we have checked. For other receptors, HUMAN comes in 43th (AVPR1), 54th (DRD1), and 46th (OXTR) lowest places. Species level analysis showed that HUMAN had the lowest variation from randomness in DRD2. Interestingly, species order in Mean(*z*) showed low correlations for different receptor pairs.

### 4.2. Intrinsic protein disorder of receptors

Intrinsic protein disorder prediction analysis revealed a high variation in the intrinsic disorder in the C- and N-terminals of the AVPR1a and DRD1 receptors (Figure 3). In AVPR1a receptor sequences high variability was observed especially in and around the residues/residue intervals 16-21, 41, 50, 55, 286-290, 453-458. Furthermore, high variability was noticed for the DRD1 receptors encircling the residues/residue intervals 20, 30-40, 210-220, 350, 435-440, and 460. In DRD2 receptors high variability was observed in and around the residue-regions 99-123, 137-147, 156, 293, 414, and 651-653. Around the residue regions 168-180, 185-194, 201-203, 209-212, 224-228, and 619-625 of the OXTR receptors high variability of intrinsic disorder was noted.

**Figure 3:**
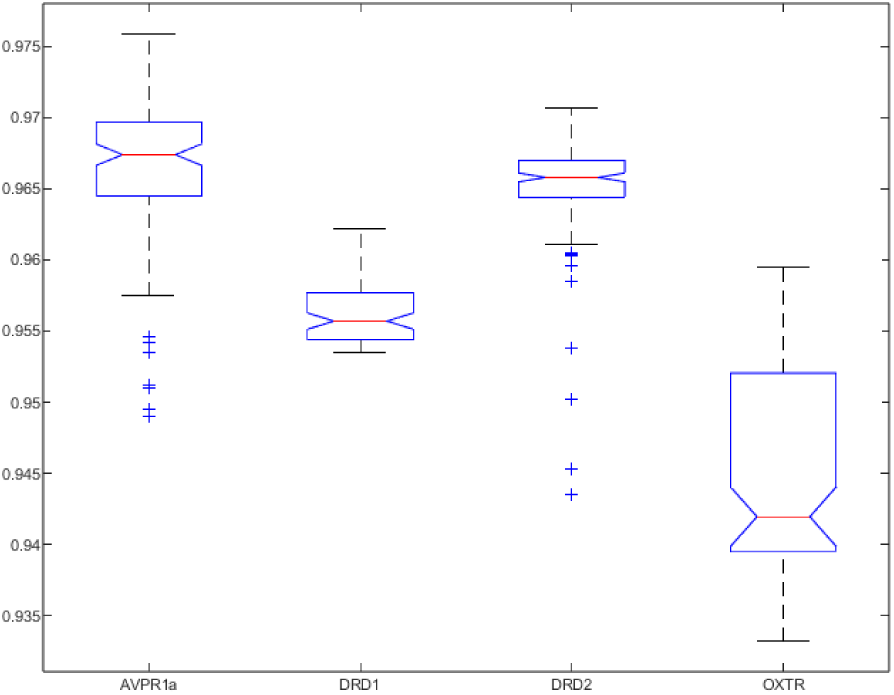
Structure and intrinsic disorder propensity of the AVPR1a, DRD1, DRD2, and OXTR receptor sequences

Disordered residues of 53.2%, 49.77%, 36.55%, and 28.08% disordered residues were found in DRD2, OXTR, AVPR1a, and DRD1, respectively. The percentages of highly flexible residues in AVPR1a, DRD1, OXTR, and DRD2 were found to be 52.68%, 47.87%, 40.28%, and 32.08% respectively. Among four different receptors, DRD2 possessed the highest number of disordered residues and the lowest amount of highly flexible residues.

### 4.3. Shannon entropy of receptors

Median Shannon entropy was noticed to be highest for the AVPR1a receptors, while lowest for OXTR (Figure 4. It is important to note that sequence diversity does not necessarily mean functional diversity. Receptors with different sequences can have similar or even identical functions, as long as their binding sites are conserved.

**Figure 4:**
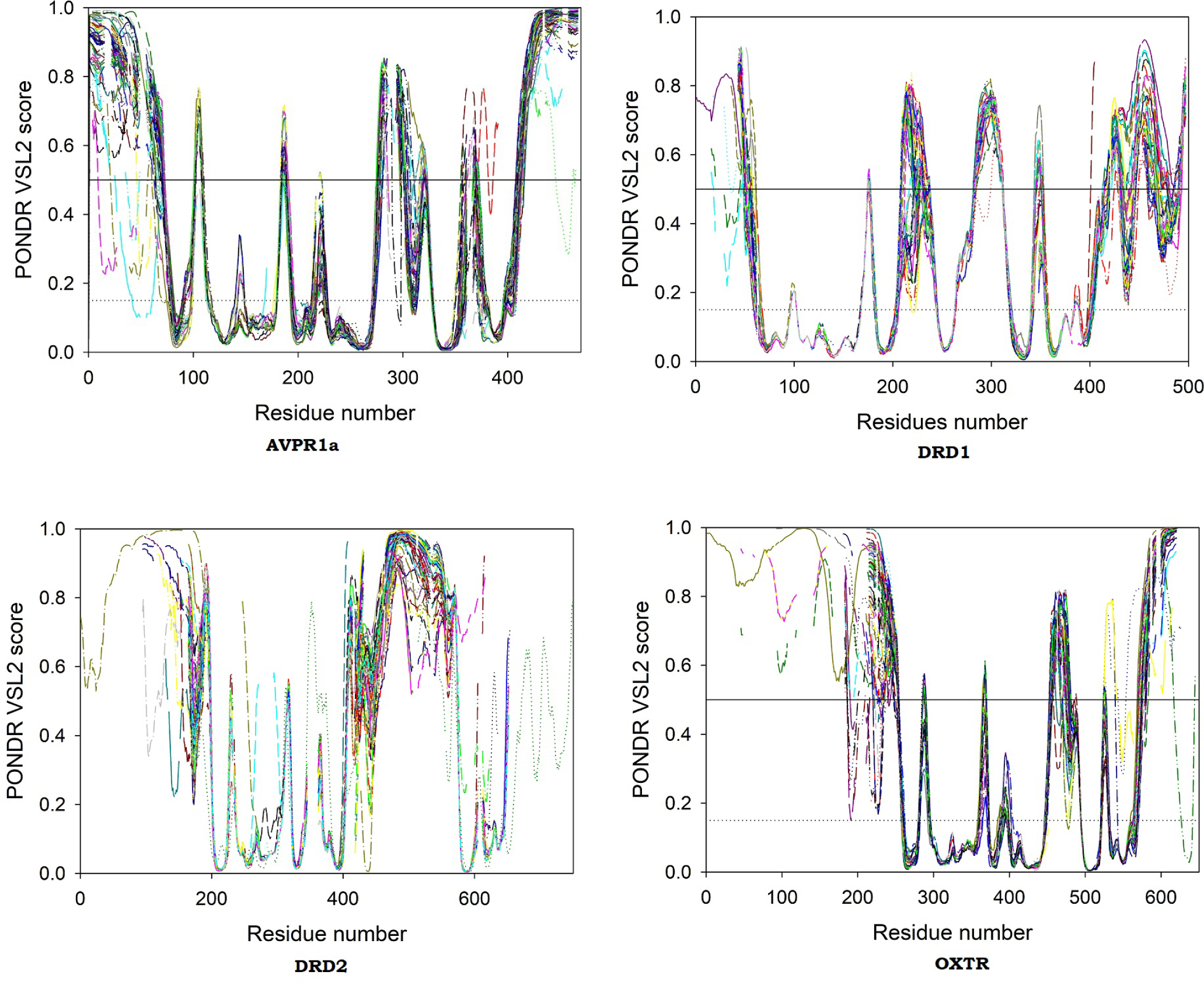
Box plot depicting variation of Shannon entropy among the four receptors

### 4.4. Shannon variability of amino acid residue positions

The lowest percentage of highly conserved amino acid residues was noticed for the AVPR1a receptors, while the highest percentage was observed for DRD1 receptors. In contrast, the highest and lowest percentages of conserved amino acids for AVPR1a and DRD1 receptors were noticed, respectively (Table 3).

**Table 3:**
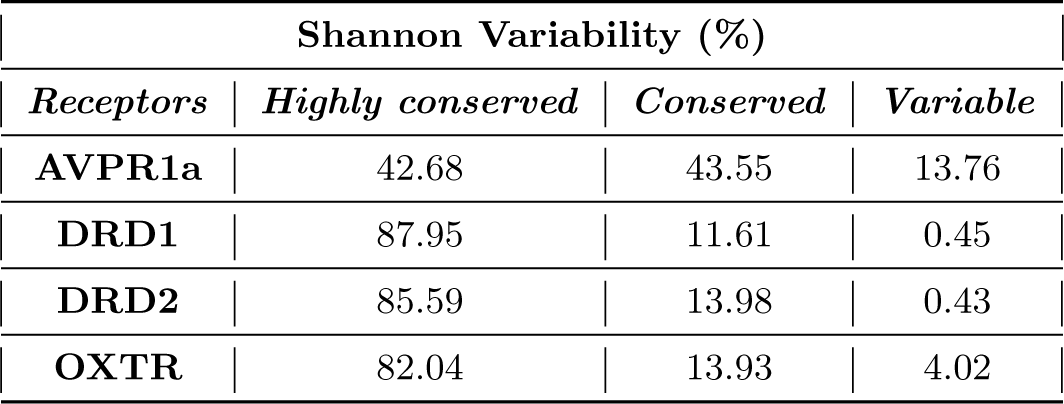
Shannon variability of amino acid residue conservation across the AVPR1a, DRD1, DRD2, and OXTR receptor sequences.

| Shannon Variability (%) |  |  |  |
| --- | --- | --- | --- |
| Receptors | Highly conserved | Conserved | Variable |
| AVPR1a | 42.68 | 43.55 | 13.76 |
| DRD1 | 87.95 | 11.61 | 0.45 |
| DRD2 | 85.59 | 13.98 | 0.43 |
| OXTR | 82.04 | 13.93 | 4.02 |

The highest sparsity with regards to the amino acid positions of receptors was noticed in DRD1, whereas the lowest sparsity was observed for the AVPR1a receptors (Figure 5).

**Figure 5:**
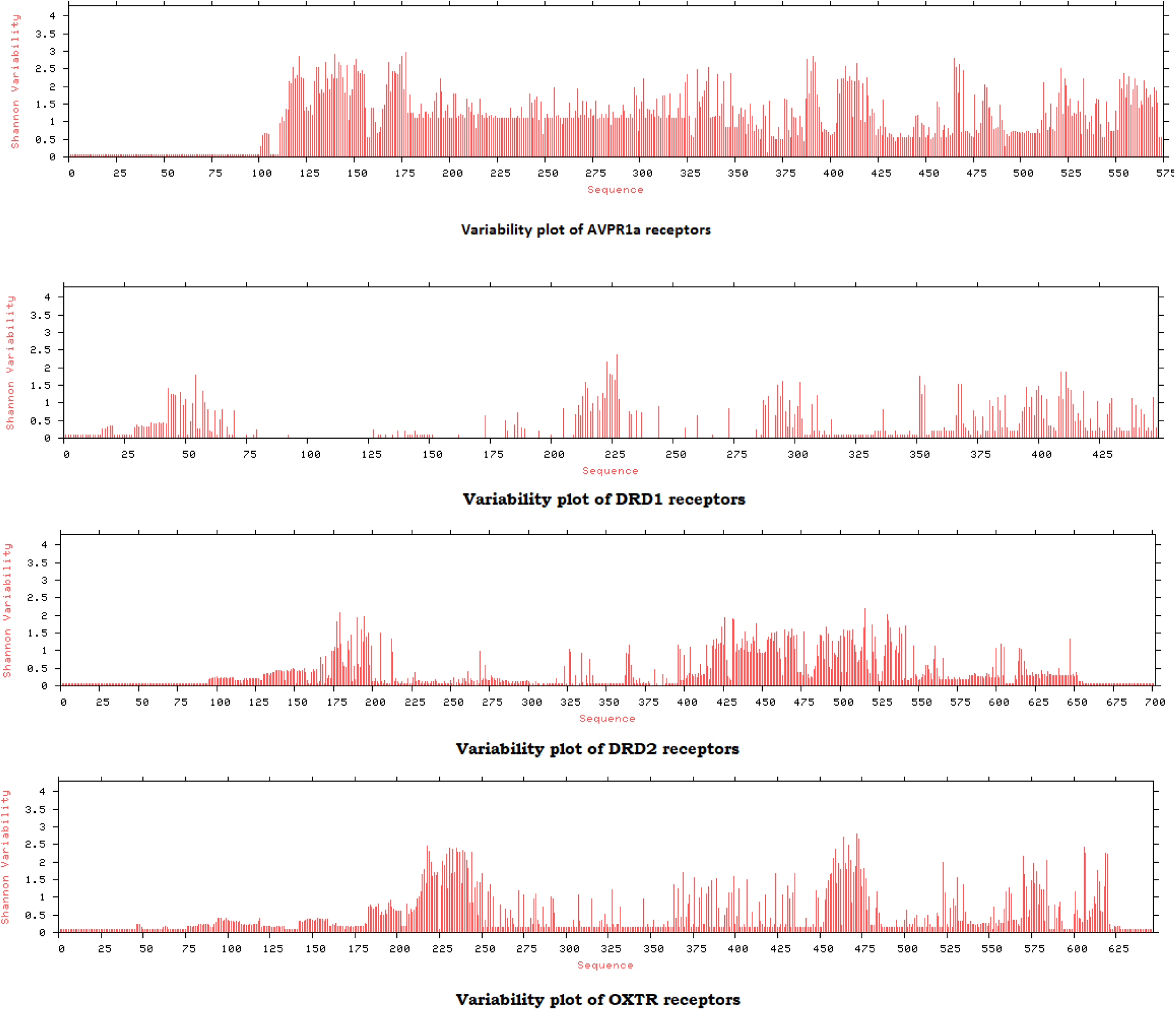
Shannon variability of amino acid residue conservation across the AVPR1a, DRD1, DRD2, and OXTR receptor sequences.

### 4.5. Binding affinity of receptors and their phylogeny

Table 4 and Figure 6 characterize the data from the docking results. Notably, DRD1 and DRD2 distributions are quite similar, and AVPR1a has higher variability than the rest. Outlier receptors tended to come from a single receptor type in species with multiple receptors recorded, such as a DRD2 from AQUCH A0A663E6H7, an OXTR from MOUSE A0A0N4SVY6, and several AVPR1a receptors across multiple species. OXTR from PANTR A0A2I3SEE8 was unable to be run as the UniProt sequence was too short to generate a valid homology model.

**Table 4:**
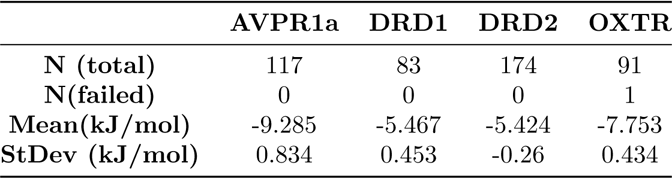
Binding affinity of four receptors

**Figure 6:**
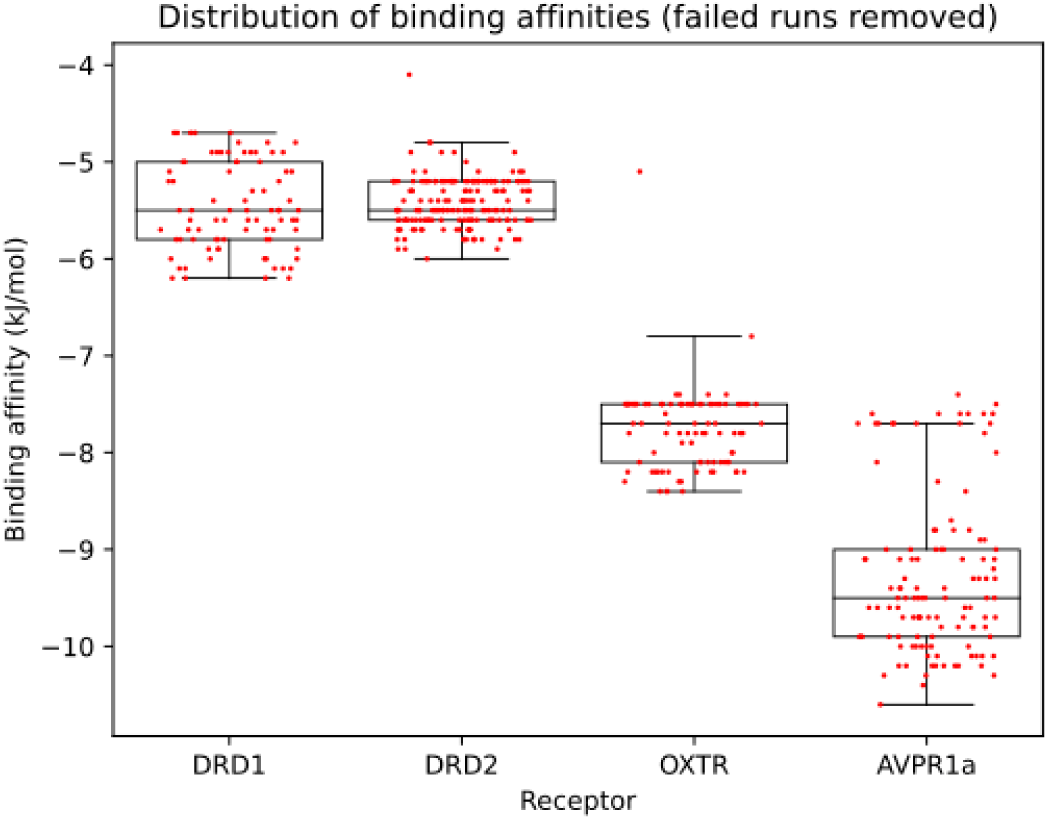
Distribution of binding affinity

Figures (7–10) show these affinities normalized with respect to each receptor (on an RGB heatmap) and arranged phylogenetically. Blue represents lower affinity with a higher kJ/mol binding energy while red represents the opposite, and green is in the middle. Each black node represents a phylogenetic split as recorded in the NCBI database, and so the relative evolutionary distances can be counted. Basic qualitative conclusions can be derived from the closeness of phylogenies and color. The weak affinity outlier pattern is much more visible as streaks of blue in DRD2, OXTR, and AVPR1a. Notably, the main outlier in terms of having stronger binding is AVPR1a from ZALCA A0A6J2D9B7, which was worth investigating. Receptors DRD2, DRD1, and OXTR had multiple data points of comparable strong binding affinity. The strongest trend is that when considering one receptor type, receptors from the same species generally have similar affinities, although sometimes phylogenetically close species tend to have similar affinities as well (most visible in OXTR).

**Figure 7:**
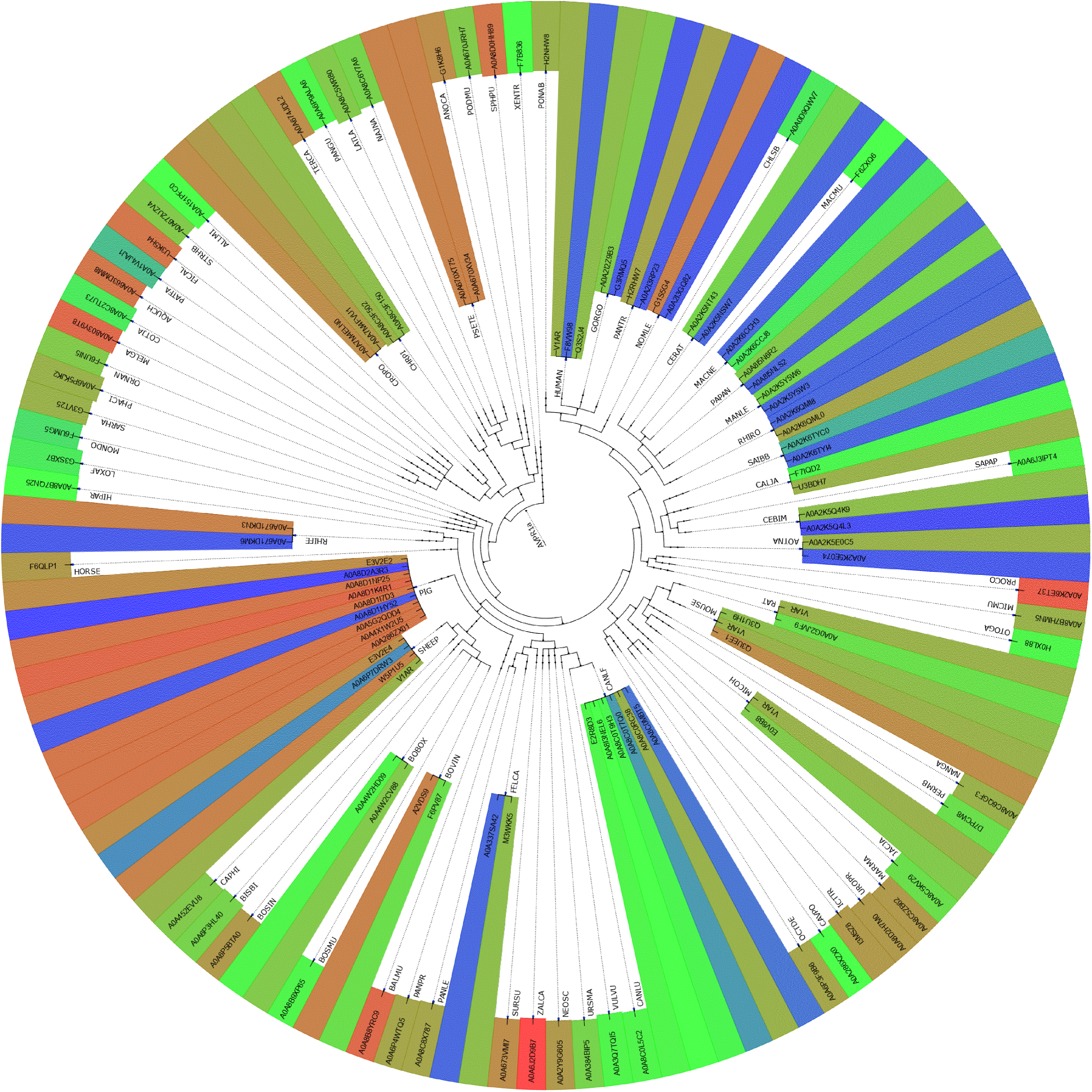
Phylogenetic relationships of AVPR1a receptor sequences based on binding affinities

**Figure 8:**
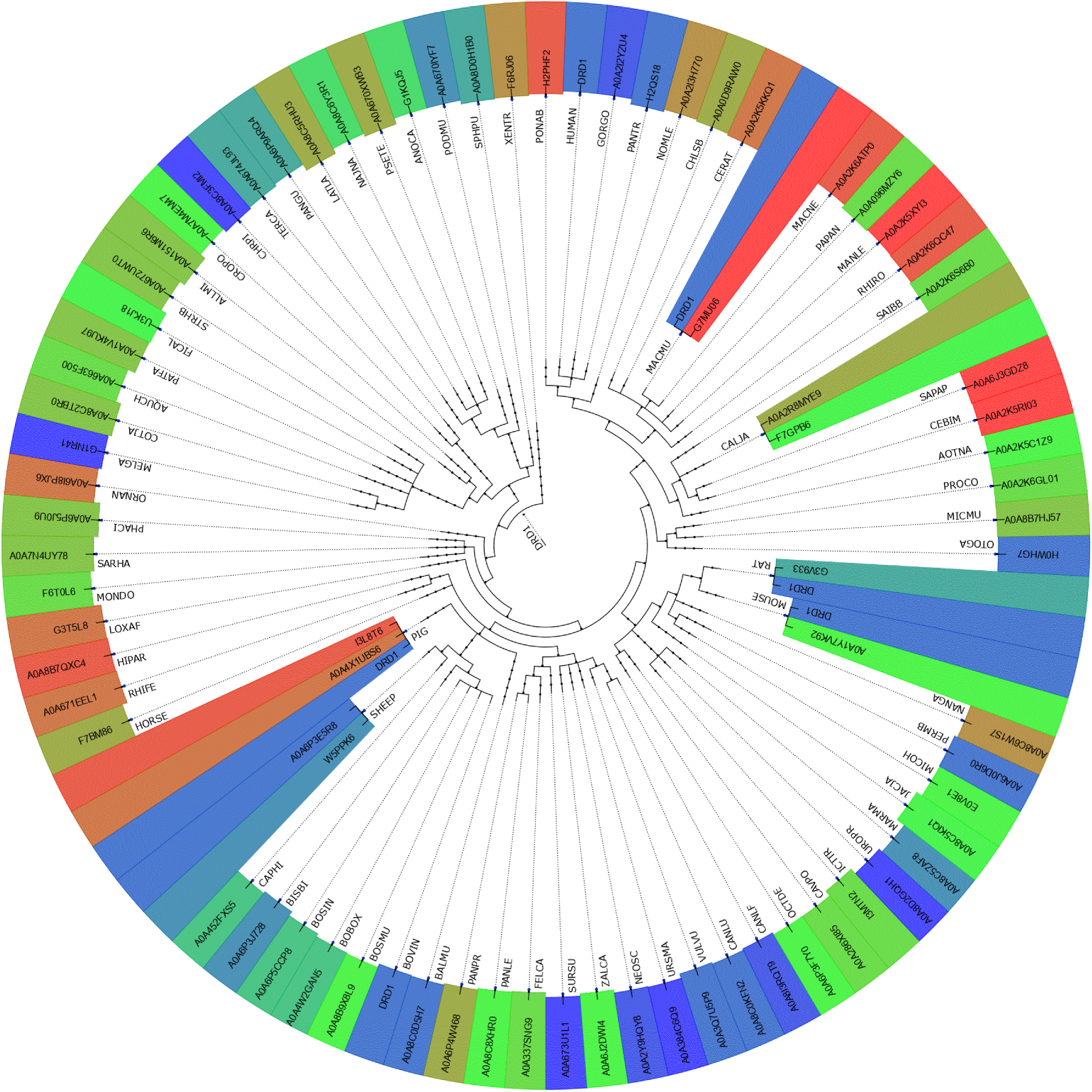
Phylogenetic relationships of DRD1 receptor sequences based on binding affinities

**Figure 9:**
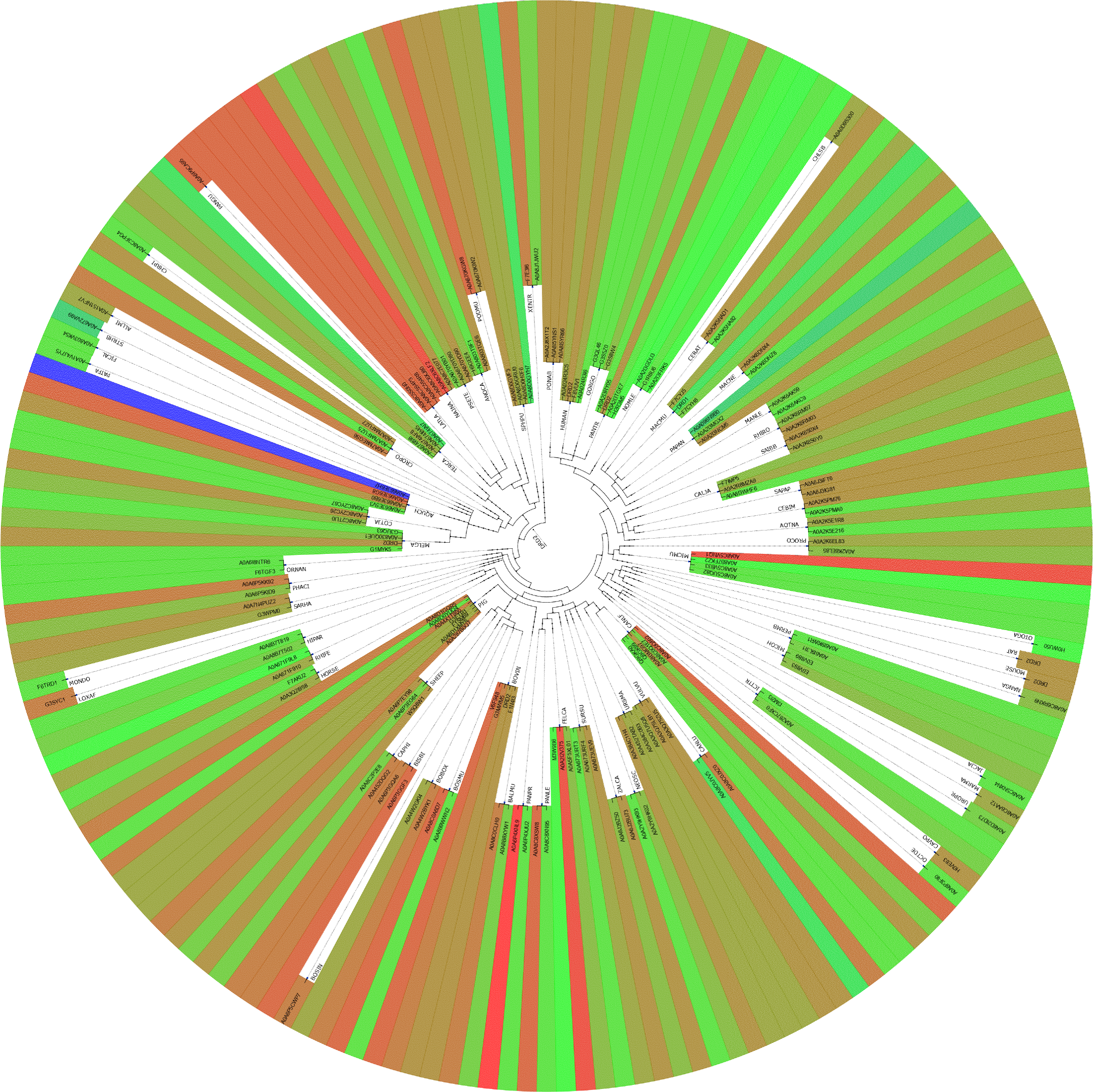
Phylogenetic relationships of DRD2 receptor sequences based on binding affinities

**Figure 10:**
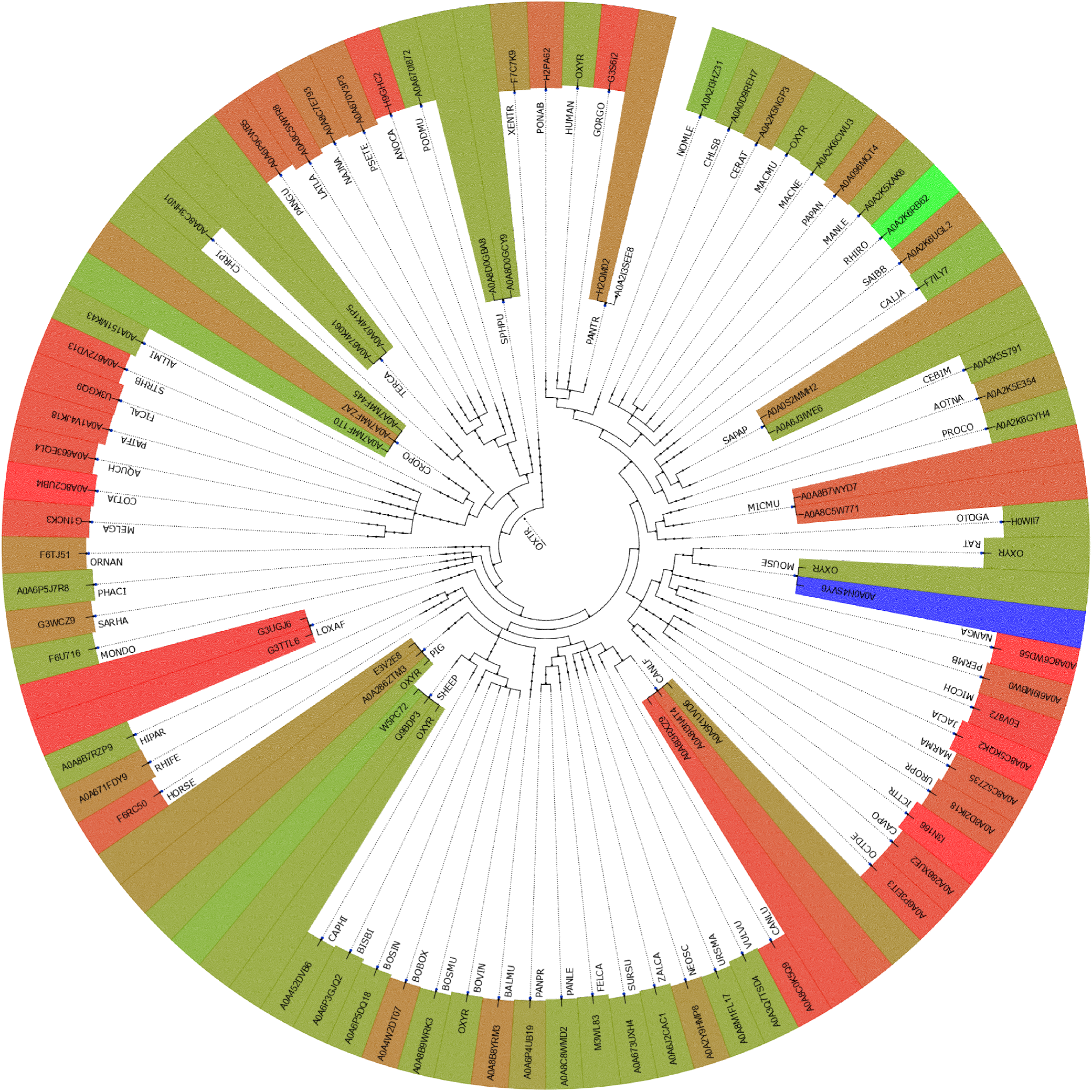
Phylogenetic relationships of OXTR receptor sequences based on binding affinities

### 4.6. Homology based SNPs and pathogenicity across receptors

A total of 24, 36, and 63 SNPs (as defined in section 3.1.6) were identified based on multiple sequence alignment of each receptor sequences of AVPR1a, DRD1, and DRD2, respectively (Table 5). It was noted that no SNPs were found for the OXTR receptor sequences. It turned out that among these four receptors, the OXTR receptor sequences were most diverged from one another.

**Table 5:** SNPs and their stability and pathogenicity

| DRD1 |  |  | DRD1 |  |  | DRD2 |  |  | AVPR1a |  |  |
| --- | --- | --- | --- | --- | --- | --- | --- | --- | --- | --- | --- |
| SNPs | Pathogenicity (PredictSNP) | Stability (iStable) | SNPs | Pathogenicity (PredictSNP) | Stability (iStable) | SNPs | Pathogenicity (PredictSNP) | Stability (iStable) | SNPs | Pathogenicity (PredictSNP) | Stability (iStable) |
| D9M | Neutral | Increase | L200V | Neutral | Decrease | S25T | Neutral | Increase | M222I | Neutral | Decrease |
| E16D | Neutral | Increase | I211L | Neutral | Decrease | D80S | Deleterious | Decrease | M222V | Neutral | Increase |
| E16K | Neutral | Decrease | S229A | Neutral | Increase | E95K | Deleterious | Decrease | T223A | Neutral | Decrease |
| E16Q | Neutral | Increase | N240S | Neutral | Decrease | C126S | Deleterious | Decrease | V226I | Neutral | Decrease |
| F21V | Neutral | Decrease | Q242H | Neutral | Decrease | I128L | Deleterious | Decrease | V228I | Neutral | Decrease |
| I23V | Neutral | Decrease | E252D | Neutral | Decrease | I128M | Deleterious | Decrease | V229A | Neutral | Decrease |
| L24V | Neutral | Decrease | Q255H | Neutral | Increase | A137S | Neutral | Decrease | V231M | Deleterious | Decrease |
| T25A | Neutral | Increase | S259N | Neutral | Increase | M140F | Neutral | Decrease | T236V | Neutral | Decrease |
| A26G | Neutral | Increase | F260L | Neutral | Decrease | N143E | Neutral | Increase | H244N | Neutral | Decrease |
| L31V | Neutral | Decrease | V280A | Neutral | Decrease | R145K | Neutral | Decrease | H244R | Neutral | Increase |
| L34I | Neutral | Decrease | I290V | Neutral | Decrease | S147T | Neutral | Decrease | I245L | Neutral | Decrease |
| L34M | Deleterious | Decrease | M294I | Neutral | Decrease | K149R | Neutral | Decrease | N248K | Neutral | Decrease |
| S35T | Neutral | Decrease | V295I | Neutral | Decrease | V152A | Neutral | Decrease | N248S | Neutral | Decrease |
| I48V | Neutral | Decrease | V295L | Neutral | Decrease | M155L | Neutral | Increase | S280D | Deleterious | Increase |
| A83S | Neutral | Decrease | Q304E | Neutral | Increase | A157T | Neutral | Decrease | S280N | Neutral | Increase |
| I85V | Neutral | Decrease | Q304K | Neutral | Increase | V159C | Deleterious | Decrease | V281I | Neutral | Decrease |
| A86I | Neutral | Increase | D309N | Neutral | Decrease | F164V | Neutral | Decrease | V281L | Neutral | Decrease |
| F88Y | Neutral | Increase | D309S | Neutral | Decrease | I166V | Neutral | Decrease | L304I | Neutral | Decrease |
| F91L | Neutral | Decrease | S310A | Neutral | Decrease | F172L | Neutral | Increase | L304V | Neutral | Decrease |
| N96T | Neutral | Decrease | D314N | Neutral | Decrease | G173S | Deleterious | Decrease | F310I | Neutral | Decrease |
| V118M | Neutral | Decrease | V317I | Neutral | Decrease | N175H | Deleterious | Decrease | F310L | Deleterious | Decrease |
| Q129R | Neutral | Increase | V317M | Neutral | Decrease | N176E | Neutral | Decrease | V312I | Neutral | Increase |
| K137R | Neutral | Increase | T343N | Neutral | Decrease | N176K | Neutral | Decrease | V312L | Neutral | Decrease |
| A138T | Neutral | Decrease | R349K | Neutral | Decrease | Q179K | Neutral | Increase | V312M | Deleterious | Decrease |
| A138V | Neutral | Decrease | T354A | Neutral | Decrease | Q179S | Neutral | Decrease |  |  |  |
| I141V | Neutral | Decrease | T354S | Neutral | Decrease | N186S | Neutral | Decrease |  |  |  |
| L142M | Neutral | Increase | A357V | Neutral | Increase | A188T | Neutral | Decrease |  |  |  |
| S144G | Neutral | Decrease |  |  |  | F189Y | Neutral | Decrease |  |  |  |
| V145I | Neutral | Decrease |  |  |  | V190A | Neutral | Increase |  |  |  |
| V151L | Neutral | Decrease |  |  |  | V191I | Neutral | Decrease |  |  |  |
| I156L | Neutral | Decrease |  |  |  | I195V | Neutral | Decrease |  |  |  |
| S161N | Neutral | Decrease |  |  |  | V196I | Neutral | Decrease |  |  |  |
| T176S | Neutral | Decrease |  |  |  | F202V | Neutral | Decrease |  |  |  |
| T188A | Neutral | Decrease |  |  |  | V204I | Neutral | Decrease |  |  |  |
| T188S | Neutral | Decrease |  |  |  | L207V | Neutral | Decrease |  |  |  |
| S191N | Neutral | Decrease |  |  |  | V208T | Neutral | Decrease |  |  |  |

For each SNPs mentioned, pathogenicity and stability were predicted, and the percentage of each type of SNPs was listed in Table 6.

**Table 6:** Percentage of SNPs across AVPR1a, DRD1, and DRD2 receptor sequences

| SNP |  | DRD1 | DRD2 | AVPR1a |
| --- | --- | --- | --- | --- |
| *Deleterious* | *Decrease* | 1.59 | 22.22 | 12.50 |
| *Deleterious* | *Increase* | 0.00 | 0.00 | 4.17 |
| *Neutral* | *Decrease* | 73.02 | 61.11 | 66.67 |
| *Neutral* | *Increase* | 25.40 | 16.67 | 16.67 |

Most SNPs which were detected in all three receptors were neutral and they were assumed to have no effective pathogenicity. Structural stability was predicted to be decreased in those neutral SNPs (Table 6).

### 4.7. Structural-physicochemical and protein features based hierarchical clustering

#### 4.7.1. Clustering of AVPR1a receptor sequences

Thirty and twenty nine clusters were formed from AVPR1a P-features and I-features, respectively. P-features and I-features based dendrograms, respectively, showed VULVU and PATFA were farthest from other AVPR1a sequences. In the marked phylogenies derived from I-Features and P-Features (**Supplementary file-3**), PATFA, ALLMI and XENTR were very much distant from the rest of the sequences and also were mutually distant. Eighteen clusters were identical in both P-features and I-features, while nine shared partial similarity (Figure 7). For each of the serial numbers (Sl nos.) 4 and 6 (7, 21, and 26) of Figure 11, one or more than one clusters I (clusters P) were subsets of clusters P (clusters I). Furthermore in Sl. No. 19 and 25 union of two clusters P was equivalent to clusters I. It can be noted that although JACJA was far away from ICTTR, UROPR, and MARMA in P-features, JACJA was clustered together with these three species in case of I-features (SI no. 7). Further, MELGA was adjacent to AQUCH, STRHB, COTJA and FICAL in case of P-features (SI no. 21), while MELGA was distant from above cited four sequences in I features. For Sl nos. 15 and 16, clusters P and clusters I had one or more than one AVPR1a sequences in common. One sequence of PIG (trE3V2E2E3V2E2) was clubbed together with SHEEP (Sl no. 16) for both I-features and P-features, whereas it also combined with CAPHI (Sl no. 16) in I-features. Some AVPR1a sequences of PIG were also clustered with RHIFE (Sl no. 26). Two sequences of HUMAN (sp37288V1AR, trQ3S2J4Q3S2J4) were adjacent to PANTR and PONAB for both I-features and P-features, while, another sequence of HUMAN was away from these two sequences. Two sequences of MICOH were clubbed together with that of MOUSE and PERMB in both I-feature and P-feature.

**Figure 11:**
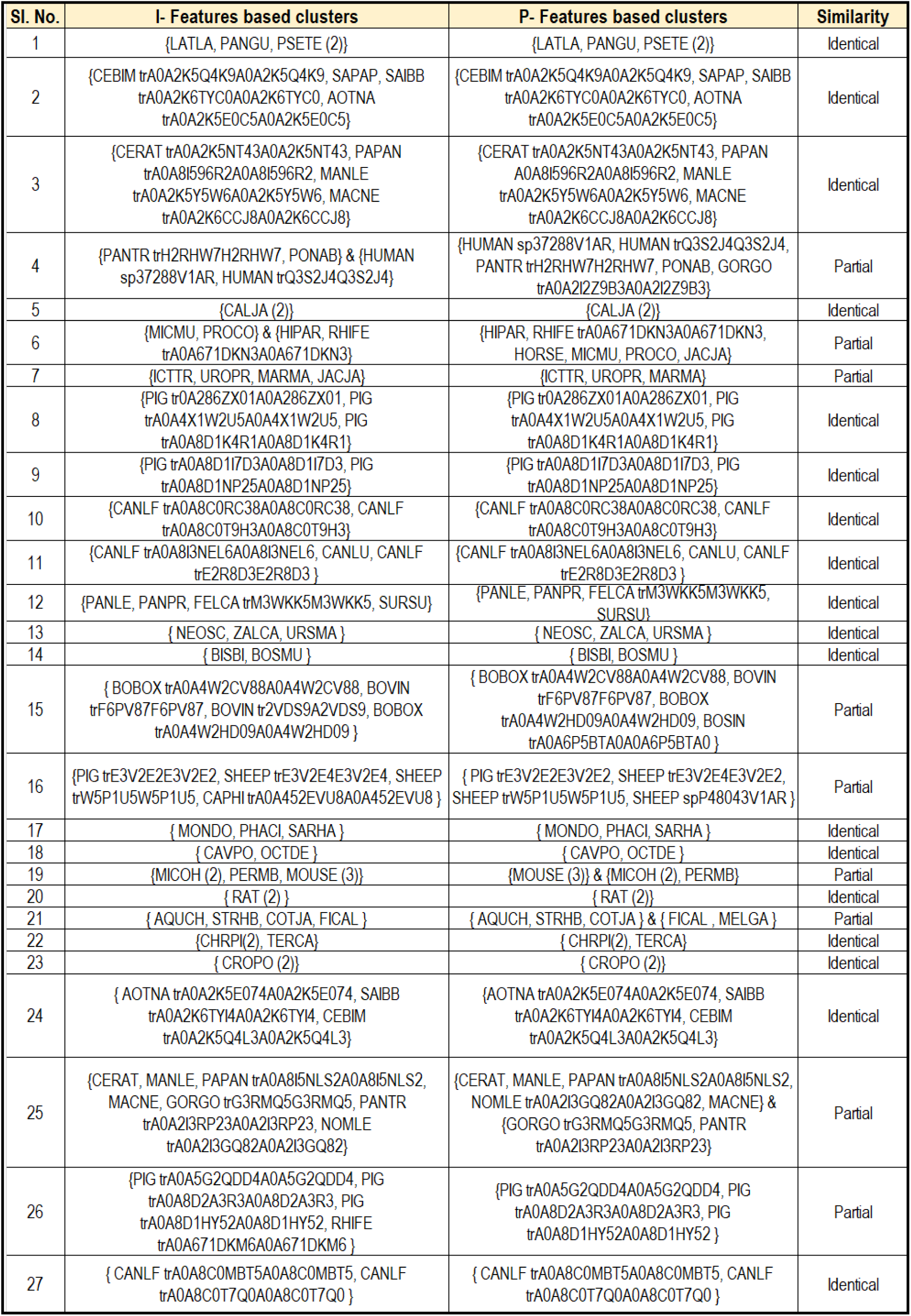
Clustering of species based on I-features and P-features of AVPR1a sequences. If two clusters formed based on I-Features and P-Features are equal then the similarity defined as ’Identical’, else, ’Partial’. SPECIES(n) signifies n AVPR1a sequences of the SPECIES belonging to the same cluster. Species tagged with Uniprot ID in a cluster signifies that particular sequence of that species belonged to the respective cluster, while other sequences of that species were placed elsewhere in AVPR1a phylogenies derived from I-Features and P-Features.

#### 4.7.2. Clustering of DRD1 receptor sequences

Twenty-one and nineteen clusters were formed from DRD1 P-features and I-features, respectively. I-features and P-features based dendrograms (**Supplementary file-3**) depicted that ALLMI, ORNAN, and XENTR were distant from all the other DRD1 sequences and also were mutually distant. In case of P-(I-)features PATFA (ANOCA) was also distant from rest of the sequences but not in case of I-(P-) features. Eight clusters were identical, while eleven clusters had partial similarity (Figure 12). In Sl nos. 1, 6, 13, and 17 clusters P were subsets of clusters I, whereas, in Sl. No. 8 and 18 clusters I were subsets of clusters P. In Sl. nos. 7, 12, 16 clusters P and clusters I had one or more than one common species. Furthermore in Sl. no. 14 union of two clusters P was equivalent to union of two clusters I. It was noteworthy that AOTNA was close to CEBIM and SAPAP in I-features, while it was distant from these two in P-features (Sl. no. 1, Figure 12). Unlike I-features CROPO was distant from TERCA and CHRPI in P-features but was close to AQUCH, MELGA, and COTJA (Sl. no. 17, Figure 12). In I-features, SPHPU was distant from NAJNA, PSETE, LATLA, and PANGU, unlike in P-features (Sl. no. 18, Figure 12). In case of P-features ANOCA and PODMU were clubbed together, whereas, they were far apart in I-features (Sl.no. 19, Figure 12). HUMAN was closest to GORGO and PANTR for both P- and I-features. MICOH was immediately adjacent to PERMB in both P-features and I-features.

**Figure 12:**
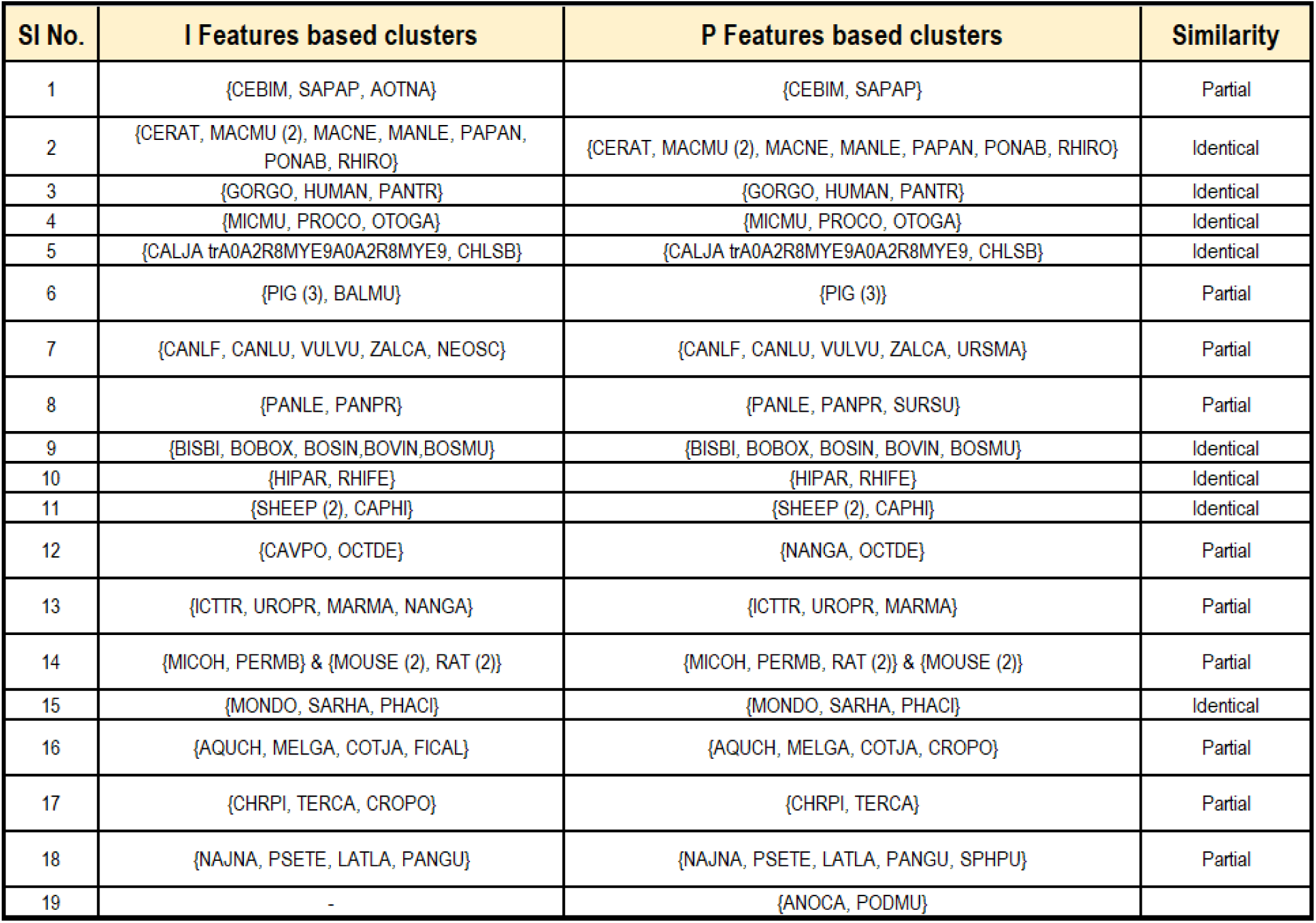
Clustering of species based on I-features and P-features of DRD1 sequences. If two clusters formed based on I-Features and P-Features are equal then the similarity defined as ’Identical’, else, ’Partial’. SPECIES(n) signifies n DRD1 sequences of the SPECIES belonging to the same cluster. Species tagged with Uniprot ID in a cluster signifies that particular sequence of that species belonged to the respective cluster, while other sequences of that species were placed elsewhere in DRD1 phylogenies derived from I-Features and P-Features.

#### 4.7.3. Clustering of DRD2 receptor sequences

Forty-four and fifty clusters were formed from DRD2 I-features and P-features (**Supplementary file-3**), respectively, among which twenty-four clusters were identical. In Sl nos. 10, 17, and 35 (16, 23) one or more than one clusters P (clusters I) were sub-clusters of clusters I (clusters P) (Figures 13 and 14). One or more than one species was/were common between clusters I and cluster P in Sl. nos. 11, 27, and 32. Furthermore in Sl. nos. 14, 15, 21, 26, 29, and 31 union of clusters P was equivalent to union of clusters I. Both the sequences of MICOH were adjacent to PERMB (trA0A6I9L3I1A0A6I9L3I1), RAT, and MOUSE in I-features and P-features (Figures 13 and 14). Unlike I-features, NANGA was close to MICOH in P-features. In both I-features and P-features HUMAN was close to GORGO, PANTR, PONAB, and RHIRO.

**Figure 13:**
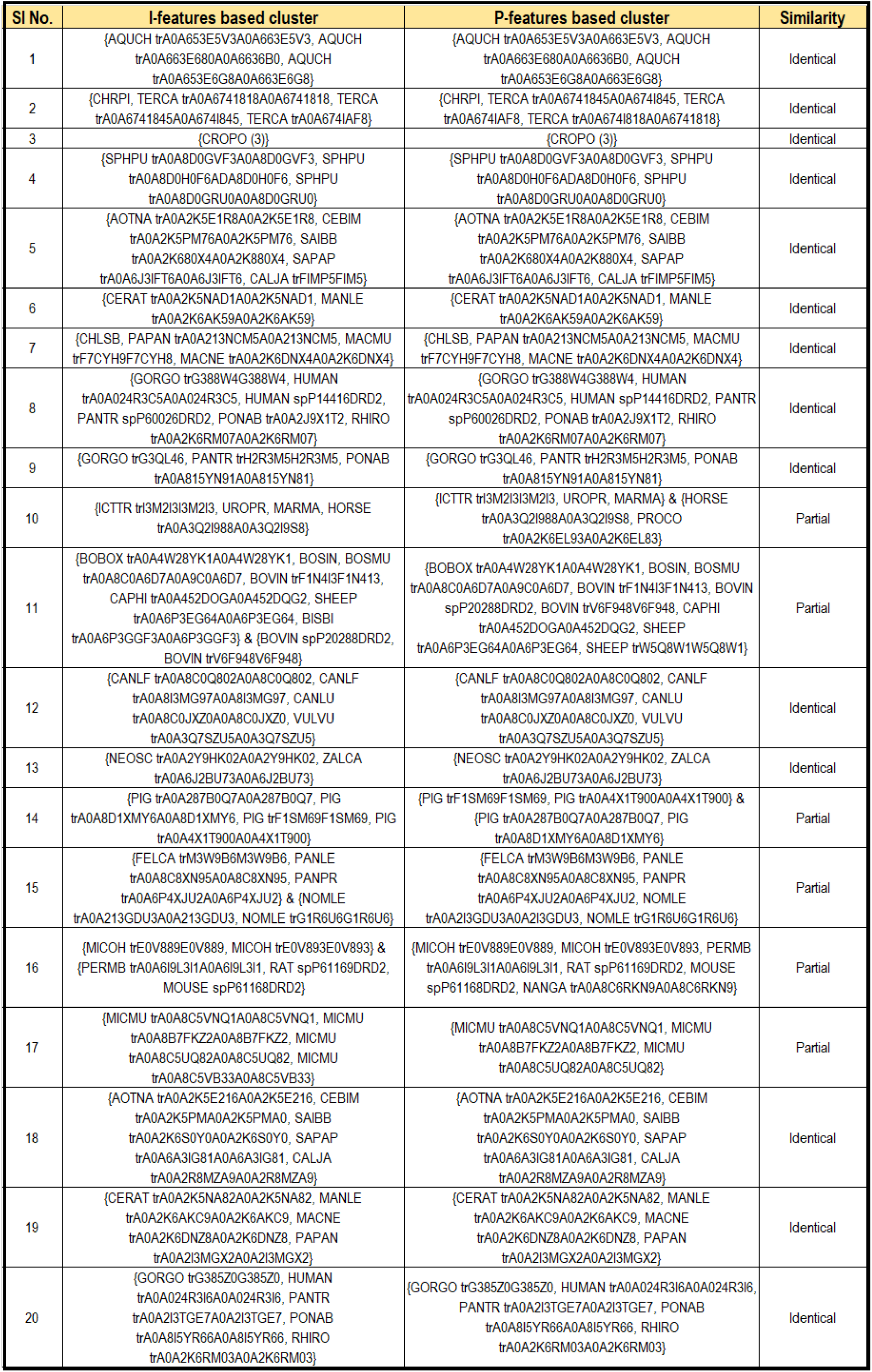
Clustering of species based on I-features and P-features of DRD2 sequences. If two clusters formed based on I-Features and P-Features are equal then the similarity defined as ’Identical’, else, ’Partial’.SPECIES(n) signifies n DRD2 sequences of the SPECIES belonging to the same cluster. Species tagged with Uniprot ID in a cluster signifies that particular sequence of that species belonged to the respective cluster, while other sequences of that species were placed elsewhere in DRD2 phylogenies derived from I-Features and P-Features.

**Figure 14:**
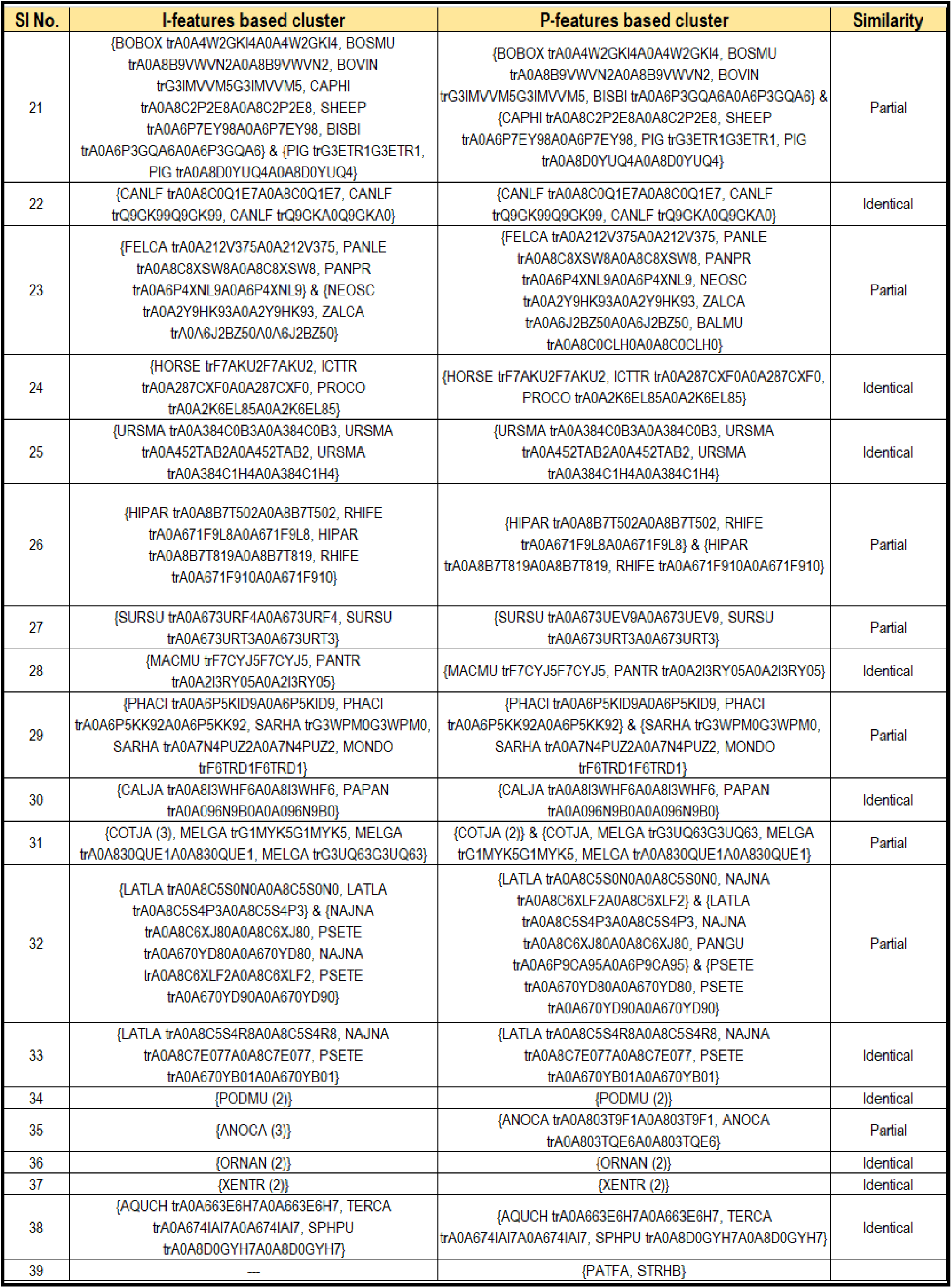
Clustering of species based on I-features and P-features of DRD2 sequences. If two clusters formed based on I-Features and P-Features are equal then the similarity defined as ’Identical’, else, ’Partial’. SPECIES(n) signifies n DRD2 sequences of the SPECIES belonging to the same cluster. Species tagged with Uniprot ID in a cluster signifies that particular sequence of that species belonged to the respective cluster, while other sequences of that species were placed elsewhere in DRD2 phylogenies derived from I-Features and P-Features.

#### 4.7.4. Clustering of OXTR receptor sequences

Twenty clusters were formed from both OXTR I-features and P-features . Both I-features and P-features based dendrograms (**Supplementary file-3**) depicted PANTR (trA0A2I3SEE8A0A213SEE8), MOUSE (trA0A0N4SVY6A0A0N4SVY6), and MICMU (trA0A8B7WYD7A0A8B7WYD7) were most distant from the other OXTR sequences. In case of I–features XENTR and MELGA were also distant from the rest of the OXTR sequences, but not in the case of P–features. Again, CROPO (trA0A7M4F445A0A7M4F445) and STRHB were distant from the rest of the sequences based on P-features, but not based on I–features. Eleven clusters were partially similar to each other, while six were identical (Table). In Sl nos. 1, 2, 3, 7, and 14 clusters P were subset of clusters I, whereas, in SI nos. 6, 15, and 17 clusters I were subsets of clusters P (Figure 15). In Sl no. 2 STRHB was adjacent to FICAL, PATFA, and COTJA in I-features, but was distant from the above-mentioned species in P features. In Sl no. 17 of P–features, two sequences of SPHPU were immediately next to PODMU, whereas in I–features two sequences of SPHPU formed a cluster together, but PODMU was fairly distant from this cluster. In SI nos. 9, 10, and 12 clusters I and clusters P had one or more than one common species. Three OXTR sequences of PIG (spP32306OXYR, trE3V2E8E3V2E8, trA0A286ZTM3A0A286ZTM3) were nearest to each other as derived from I features (Sl no. 7), whereas one sequence (trA0A286ZTM3A0A286ZTM3) was away from the other two in case of P-features. HUMAN was close to GORGO, PANTR (trH2QM02H2QM02), PONAB and NOMLE in both I-features and P-features (Figure). For P-features MICOH was closest to HORSE, whereas in I-features it was closest to PERMB.

**Figure 15:**
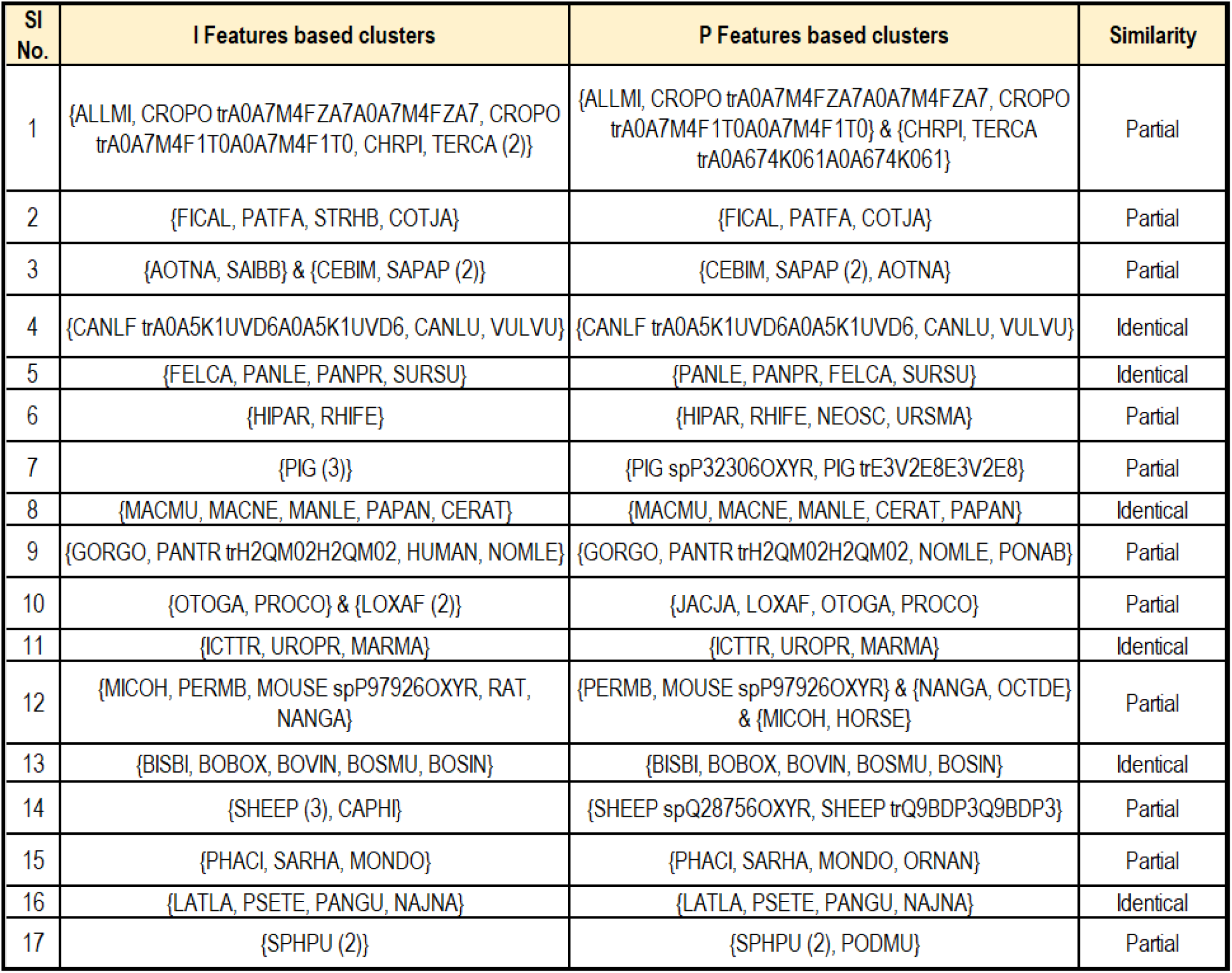
Clustering of species based on I-features and P-features of OXTR sequences. If two clusters formed based on I-Features and P-Features are equal then the similarity defined as ’Identical’, else, ’Partial’. SPECIES(n) signifies n OXTR sequences of the SPECIES belonging to the same cluster. Species tagged with Uniprot ID in a cluster signifies that particular sequence of that species belonged to the respective cluster, while other sequences of that species were placed elsewhere in OXTR phylogenies derived from I-Features and P-Features.

### 4.8. Proximal relationships among species based on AVPR1a, DRD1, DRD2, and OXTR receptors

Quantitative features of the four receptors AVPR1a, DRD1, DRD2, and OXTR as extracted from the respective protein sequences led to a set of proximal relationships among various species. Species (receptor sequence of species) are proximally related, if they belong to the Intersection of clusters as derived from I-features and P-features of a receptor.

LATLA, PANGU, and PSETE were related as they all belong to the intersection of clusters derived from I-features and P-features in the case of AVPR1a (Sl. No. 1 (Figure 11). Likewise, HUMAN, PANTR, and PONAB were related in AVPR1a (Sl. No 4 (Figure 3)). Note that, GORGO was not related with HUMAN, PANTR, and PONAB as it does not appear in the intersection of the clusters (Sl. No 4 (Figure 11)). In a similar fashion, relationships among species were derived in the cases of all four receptors.

We considered clusters that were present in two or three out of four receptors. The clusters [*{*GORGO, HUMAN, PANTR*}*, *{*MICOH, PERMB, MOUSE, RAT*}*, *{*COTJA, MELGA*}*] were common among DRD1 and DRD2. [*{*MICMU, PROCO*}*, *{*AQUCH, COTJA*}*] clusters were present in AVPR1a and DRD1. [*{*NEOSC, ZALCA*}*, *{*CEBIM, SAPAP, SAIBB, AOTNA*}*, *{*PANTR, PONAB*}*] clusters were common in AVPR1a and DRD2. [*{*GORGO, PANTR, NOMLE*}*, *{*MACMU, MACNE, MANLE, PAPAN, CERAT*}*, *{*PANLE, PANPR, FELCA, SURSU*}*, *{*COTJA, FICAL*}*] were common in AVPR1a and OXTR. The clusters [*{*OTOGA, PROCO*}*, *{*MACMU, MANLE, PAPAN*}*] were common in OXTR and DRD1.

The cluster *{*LATLA, PANGU, PSETE*}* was present in AVPR1a, DRD1, and OXTR. [*{*CEBIM, SAPAP, AOTNA*}*, *{*PANLE, PANPR, FELCA*}*, *{*PIG, SHEEP*}*] clusters were common in AVPR1a, OXTR, and DRD2. [*{*CANLF, CANLU, VULVU*}*, *{*BISBI, BOBOX, BOVIN, BOSMU, BOSIN*}*, *{*HIPAR, RHIFE*}*, *{*LATLA, PSETE, NAJNA*}*, *{*PAPAN, MACMU*}*] these discrete clusters were common among OXTR, DRD1, and DRD2.

Considering all four receptors AVPR1a, DRD1, DRD2, and OXTR, the common clusters among all species were taken into account (Figure 16). Likewise, eleven discrete clusters were formed with twenty-nine different species (Figure 17, 18). Generally, species belonging to the same order were clubbed together and their behavioural pattern in terms of polygamy and monogamy were found to be similar. This depicts that in general the genetic patterns are translationally related to the behavioural patterns. Conversely, some species belonging to the same order but having dissimilar behavioural pattern were found to fall in the same cluster based on I-features and P-features. For instance, although ICTTR, UROPR, and MARMA belonged to the same phylum-class and order, ICTTR and UROPR showed polygamous nature, whereas MARMA was monogamous (Figure 2). Again, CANLF and CANLU belonged to same order and formed a single cluster, although CANLF is polygamous and CANLU is monogamous. MOUSE, MICOH, and PERMB belonging to the same order formed a single cluster, but with different coital behaviour.

**Figure 16:**
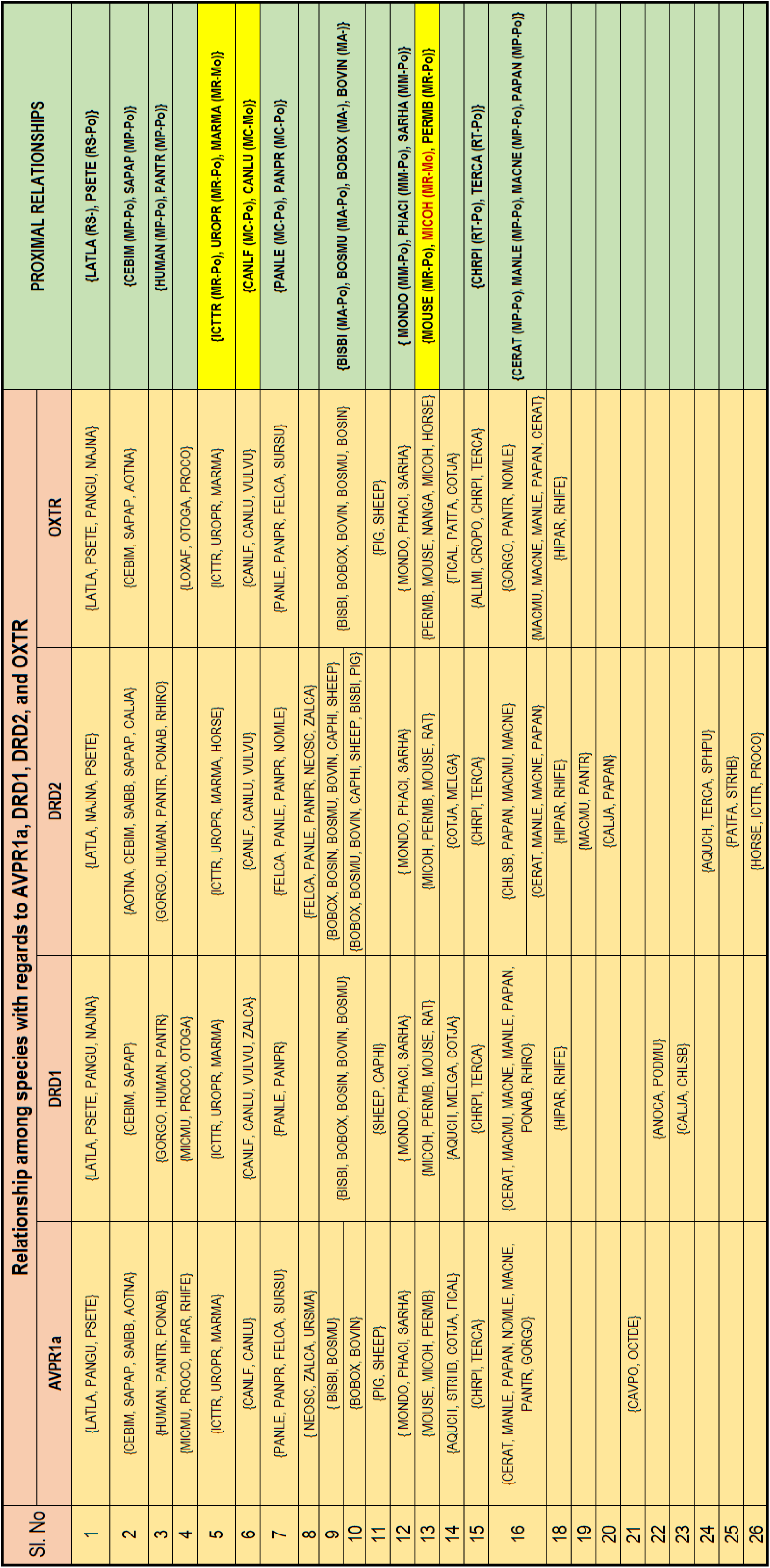
29 proximally related species based on I-features and P-features of the four receptors. Mo-Monogamous, Po-Polygamous. Highlighted in yellow represents the set of proximally related species which showed different coital behavior.

**Figure 17:**
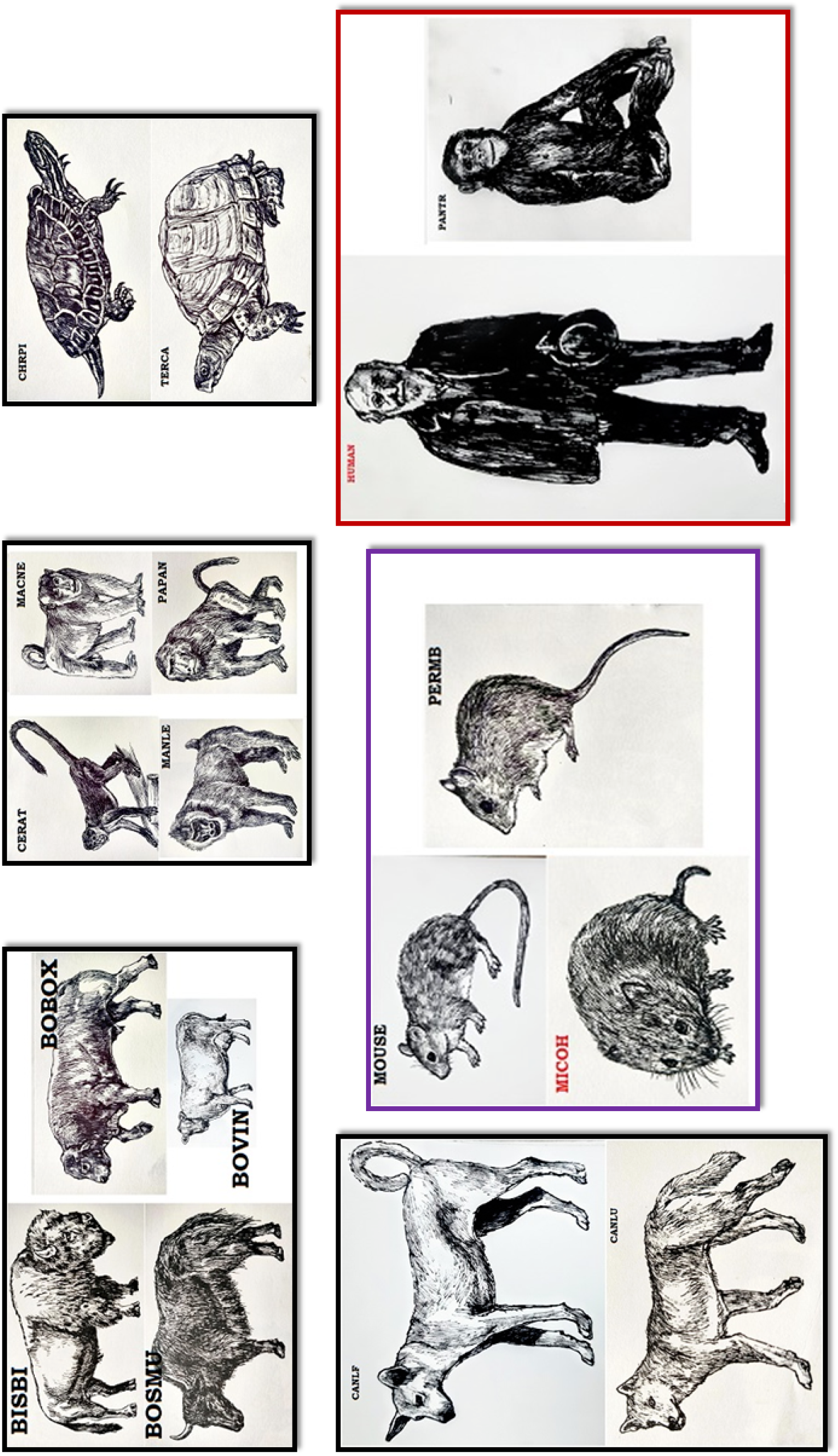
Proximal relationships among species

**Figure 18:**
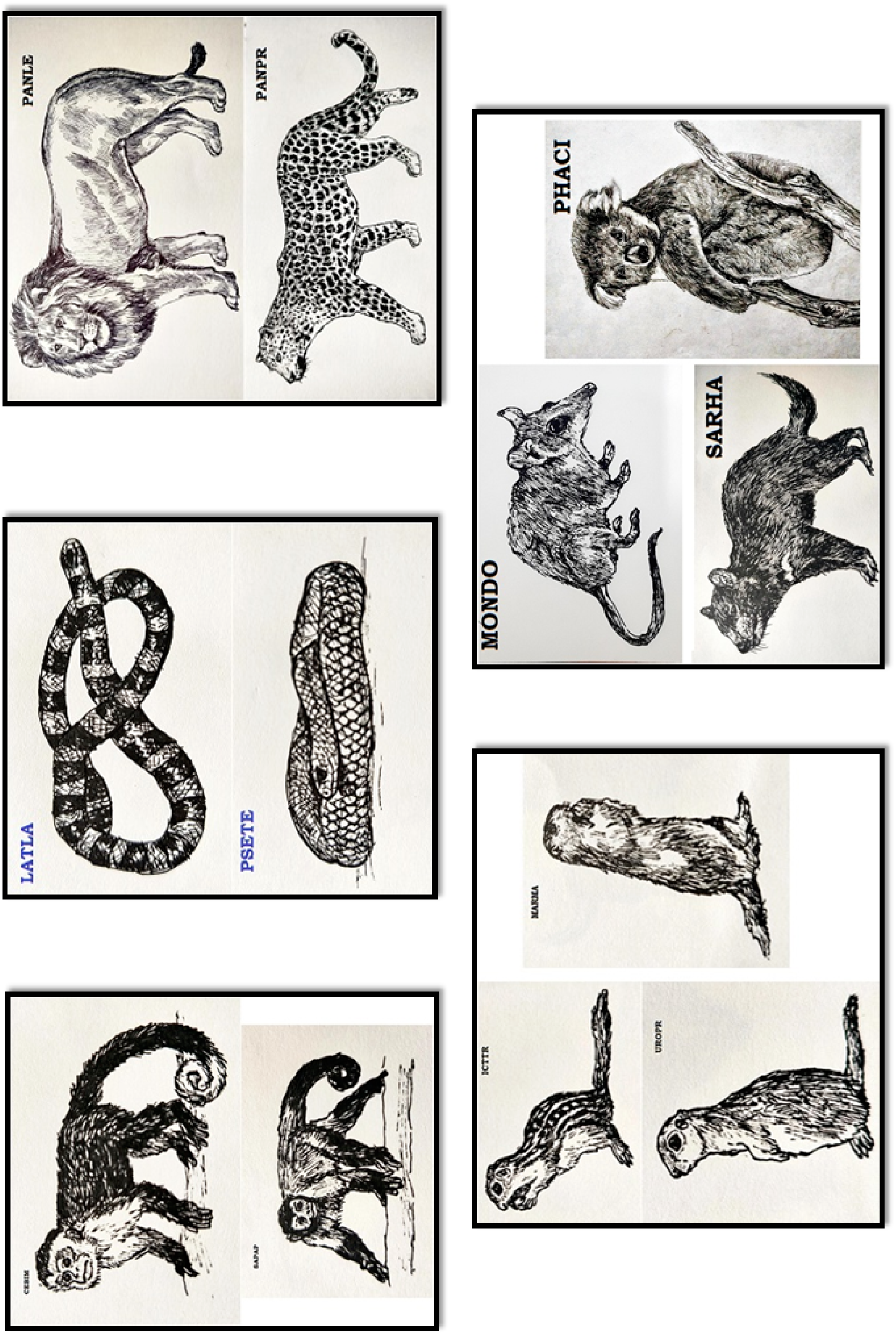
Proximal relationships among species

HUMAN and MICOH (prairie vole) belonged to different clusters for each receptor. HUMAN was closest to PANTR while MICOH was closest to MOUSE and PERMB when four receptor sequences were taken into account.

Furthermore, it was noted that all the proximally related species were phylogenetically close to each other based on binding affinities of all four receptors, while MOUSE, MICOH, and PERMB were placed a little further away from each other in each phylogeny (Figures 7–10).

## 5. Discussions and concluding remarks

Quantitative analyses of receptors showed that AVPR1a possessed the highest median of Shannon entropy. Highest percentage of conserved amino acid and lowest sparsity in the variability plot was found in AVPR1a receptors as obtained from Shannon variability analysis. On contrast, the lowest percentage of conserved amino acid and the highest sparsity in the variability plot was noticed in DRD1 receptors. The lowest deviation from randomness based on polarity of amino acid sequences and the lowest intrinsic disordered residue of the receptors were found in case of DRD1 receptors. The highest deviation from randomness based on polarity was found in the case of DRD2. Interestingly it was further noted that among all species, humans showed the highest randomness in case of DRD2. Also, DRD2 had the highest number of disordered residues and the lowest number of flexible residues as obtained from Intrinsic protein disorder analysis. the highest number (63) of homology-based SNPs was seen for the DRD2 receptors, depicting the lowest diversity among the residual positions of the aligned sequences. The median Shannon entropy was the lowest for OXTR, portraying skewed frequency distribution of amino acid sequences. No SNP was found for OXTR receptor sequences which characterized the sequence as highly diverse.

Altogether quantitative analyses of these four receptor sequences revealed that AVPR1a, DRD1, DRD2, and OXTR were distinctively different from each other and their cumulative contributions govern the coital behavior of various species.

LATLA, PANGU, and PSETE were found to be proximally related species, showing proximity of coital behaviour. Similar results were found in the case of another 7 sets of proximally related species (Figure 16). Dobzhansky thoroughly examined the link between genetic makeup and behavioral characteristics. Although nature is not inherently deterministic, our physical attributes and resulting traits are essentially byproducts of gene replication [97]. On the contrary, although MOUSE, MICOH, and PERMB were proximally related, they showed different mating behaviour. In previous study it has been reported that cuckoo inspite of being genetically polygamous showed social monogamy possibly due to its nature of brood parasitism [98]. Thus, from here we can certainly infer that species having closest proximal relation in terms of genetic makeup differs in their behavioural pattern, mostly, due to behavioural epigenetic regulations. Behavioural epigenetic regulation relates to the influence of nurture on the nature of the species concerned. Here nature means biological heredity, whereas nurture means the effect of natural phenomenon introduced into their lifespan. This study speaks for the everlasting nature-nurture debate [99]. Based on I-features and P-features HUMAN and MICOH (prairie vole) are phylogenetically distant, depicting that the genetic attribute is different among these two species, also, their behavioural pattern showed a direct translation of their genetic constituents. Societal norms (factors inducing epigenetics) though impose us to become monogamous, genetically humans are polygamous. Thus, although based on social, traditional, religious, ethical, and economic grounds humans have aimed for monogamous behavior for a long time, during recent decades there has been a shift towards a more polygamous society.

However, further research is needed to fully understand the mechanisms underlying behavioral epigenetic regulations and their impact on mating behavior. Additionally, exploring the genetic differences between species with similar mating behaviors could help shed light on the nature-nurture debate and the role of societal norms in shaping behavior. Overall, the present study opens up new avenues for future research on the genetic and epigenetic mechanisms underlying coital behavior.

## Acknowledgements

Authors are very grateful to the laboratory supporters Mr. Ashim Dhar and Mrs. Piyali Maitra for their logistic assistance. Authors would like to thank the Department of Biotechnology, Govt. of India (BT/PR26647/NNT/28/1365/2017), Indian Space Research Organization (ISRO/RES/3/827/19-20), and Indian Statistical Institute (ISI) (ISI/TAC/PROJECT 1/2022-23) for their financial support.

## Author contributions statement

MS and SSH conceived the problem and theoretical experiments. SSH, VNU, MS, DN, and SC executed the results and performed the analysis. DN and PB performed statistical analyses. LR and EW did the molecular docking and binding affinity derivation analysis. SSH, DN, VNU, MS, PB, KL, LR, EW, and SC wrote the initial draft. All authors reviewed and edited the manuscript. AG supervised the entire project. All the authors checked, reviewed, and approved the final version of the manuscript.

*Data Collectors*: Jacqueline He, Linh Nguyen, Srinitha Srikanth, Samarah Mohammad, Nathan Lanclos.

