## Supplementary figures and images for "Biology of Coital Behavior: Looking Through the Lens of Mathematical Genomics"

### Supplementary file-3

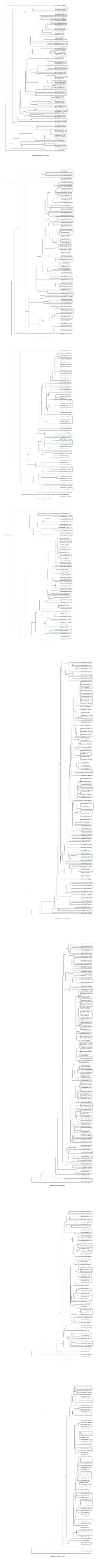
